# A neural circuit basis for context-modulation of individual locomotor behavior

**DOI:** 10.1101/797126

**Authors:** Kyobi Skutt-Kakaria, Pablo Reimers, Timothy A. Currier, Zach Werkhoven, Benjamin L. de Bivort

## Abstract

Defying the cliche that biological variation arises from differences in nature or nurture, genetically identical animals reared in the same environment exhibit striking differences in their behaviors. Innate behaviors can be surprisingly flexible, for example by exhibiting context-dependence. The intersection of behavioral individuality and context-dependence is largely unexplored, particularly at the neural circuit level. Here, we show that individual flies’ tendencies to turn left or right (locomotor handedness) changes when ambient illumination changes. This change is itself a stable individual behavioral characteristic. Silencing output neurons of the central complex (a premotor area that mediates goal-directed navigation) blocks this change. These neurons respond to light with idiosyncratic changes to their baseline calcium levels, and idiosyncratic morphological variation in their presynaptic arbors correlates with idiosyncratic sensory-context-specific turn biases. These findings provide a circuit mechanism by which individual locomotor biases arise and are modulated by sensory context.

## Introduction

Individuality, defined as persistent idiosyncratic differences in behavior, has been observed in essentially every species examined, and this variability arises even when individuals share a genotype and rearing environment (Buchanan et al., 2015; Freund et al., 2013; Horstick et al., 2019; Kain et al., 2012). Individuality can arise from neural circuits that integrate continuously varying inputs to produce a discrete behavioral output, as these circuits likely employ nonlinearities that render them sensitive to idiosyncratic stochastic fluctuations (Buchanan et al., 2015; Lebovich et al., 2019). But beyond understanding that individuality is ubiquitous and expected, its biological basis is elusive.

The complexity of even “simple” invertebrate nervous systems complicates efforts to identify causal variants that give rise to idiosyncratic behavioral biases, but there are a few promising examples. Individual *Drosophila melanogaster* larvae exhibit stochastic variation in the branching pattern of 12A hemilineage interneuron projections, and this variation predicts individual differences in the timing of flight initiation (Mellert et al., 2016). In adult flies, left-right wiring asymmetry in bilaterally-projecting visual Dorsal Cluster Neurons predicts idiosyncratic deviation from direct walking between attractive visual cues (Linneweber et al., 2019). A general approach to mapping such correlates of individual bias could be to prioritize neural bottlenecks: regions of the brain in which many inputs converge onto a relatively small number of output cells. The representation of computational features contributing to behavior may be more sensitive to stochastic fluctuations in these bottlenecks than in regions of a circuit where those features are encoded across a large population of neurons and averaging can be used to diminish the effects of stochastic fluctuations. Because of its relative simplicity, correspondence of specific cells across individuals, well-mapped nervous system and unrivaled genetic toolkit, the fruit fly is an excellent model system for mapping neural substrates of individuality (and it is not surprising that early successes have been found in this species).

How flies respond to a stimulus or behave spontaneously can depend on the sensory context, and when this flexibility is adaptive, it may represent a simple form of cognition (Gorostiza, 2018). As examples: flies prioritize or de-prioritize grooming their wings, depending on whether their eyes are dirty (Seeds et al., 2014). Light is an attractive cue to flies that are capable of flying (probably because the sky is a means of escape) but aversive to flies that can only walk (Gorostiza et al., 2016). The visual presence of parasitoids induces flies to favor more alcoholic substrates for oviposition (Kacsoh et al., 2013). Freshly eclosed flies will only perch and inflate their wings if there is enough open space around them (Peabody et al., 2009). Thus, innate behaviors can exhibit striking context-dependent flexibility.

A fly locomoting on complex natural surfaces makes decisions essentially continuously. In the lab, a fly in a Y-shaped maze makes decisions every time it comes to the center of the maze. While the average fly makes right and left turns with equal probability (i.e., has no bias), individual flies often display strong biases towards the right or left (Buchanan et al., 2015). While there is no evidence of genetic determination of turn bias, the variance of these distributions is under genetic control (Ayroles et al., 2015), and genetic effects on variability may be more important than environmental effects on variability (Akhund-Zade et al., 2019). Idiosyncratic biases appear to arise in development (Ayroles et al., 2015) and are stable throughout a fly’s lifetime (Buchanan et al., 2015).

The central complex (CX) brain region is thought to be critical to controlling oriented behaviors in insects (Honkanen et al., 2019). We found previously that morphological mutants of the CX, as well as acute silencing of neural populations within the CX, were able to abruptly broaden the distribution of biases across the population, indicating that neurons within the CX at least play a role in turn bias modulation (Buchanan et al., 2015). The CX is made up of four canonical midline spanning neuropil, the Protocerebral Bridge (PB), Ellipsoid Body (EB), Fan-shaped Body (FB), and the paired Noduli (No). The recurrent circuit between the EB and PB was shown to encode heading, a key computational element in guiding oriented behaviors. As a fly turns, a spatially localized “bump” of neural activity moves through this structure to encode the heading estimate. Both visual and non-visual inputs could update this heading representation, and when the animal was not moving, the bump remains in one place via apparent ring-attractor functionality (Giraldo et al., 2018; Green et al., 2017; Kakaria and de Bivort, 2017; Kim et al., 2017; Seelig and Jayaraman, 2015). In addition to representing features useful for orientated walking behaviors, representations in the CX show context-dependence based on an animal’s state of activity (Martin et al., 2015; Weir et al., 2013), and neurons in the CX can causally modulate internal state (Donlea et al., 2018; Liu et al., 2016).

Here, we combine high throughput behavioral quantification (Buchanan et al., 2015), genetic manipulation of specific neuron populations (Hamada et al., 2008; Kitamoto, 2001), calcium imaging from a tethered walking fly (Seelig et al., 2010) and anatomical analysis (Talay et al., 2017) to uncover a circuit mechanism of individuality in a context-dependent behavior of *Drosophila melanogaster*. We find that individual animals display different locomotor biases depending on the presence or absence of ambient illumination, a phenomena we call <u>L</u>ight-<u>D</u>ependent <u>M</u>odulation of locomotor bias (LDM). Acute physiological manipulations of neuronal activity reveal that cell types in visual pathways and specific cell populations in the central complex mediate LDM. We find that a circuit element’s effect on LDM can be sensory-context specific (i.e., preferentially affecting bias in the light or dark). Perturbing neurons (“PF-LCs”) that project from the PB into the lateral portion of the Lateral Accessory Lobe (LAL) specifically altered the locomotor biases displayed in one sensory context (when the lights were on). The activity of PF-LCs showed a light-specific correlation with turning behavior, and their anatomical asymmetry predicted stable idiosyncratic turning biases. Asymmetrically altering the physiology of these neurons with optogenetics altered locomotor handedness in a directed way, demonstrating that asymmetries in these neurons are sufficient to impart locomotor asymmetries. It therefore appears that PF-LCs are a “locus of individuality,” a site where idiosyncratic circuit differences cause idiosyncratic behavioral biases.

## Results

To examine how sensory context (specifically, changes in luminance) affects individual locomotor biases, we modified instruments previously used to track flies as they made left-right locomotor choices in symmetrical Y-shaped mazes (Ayroles et al., 2015; Buchanan et al., 2015). We added a dual-channel illuminator, under computer control, so that behavioral arenas could be lit with white LEDs, infrared LEDs, or both (Fig 1A). This instrument put flies in two conditions: “light,” with flies tracked using the IR LEDs and the white LEDs on, and “dark,” with flies tracked using the IR LEDs and the white LEDs off (Fig 1B). The IR tracking LEDs were invisible to the flies, and the white LEDs provided diffuse ambient illumination. This instrument allowed us to quantify a number of locomotor features in both dark and light conditions, with high precision, for 120 individual flies simultaneously. Using four copies of this instrument, we conducted a genetic screen to assess the role of many neural circuit elements in these aspects of behavior.

**Figure 1.**
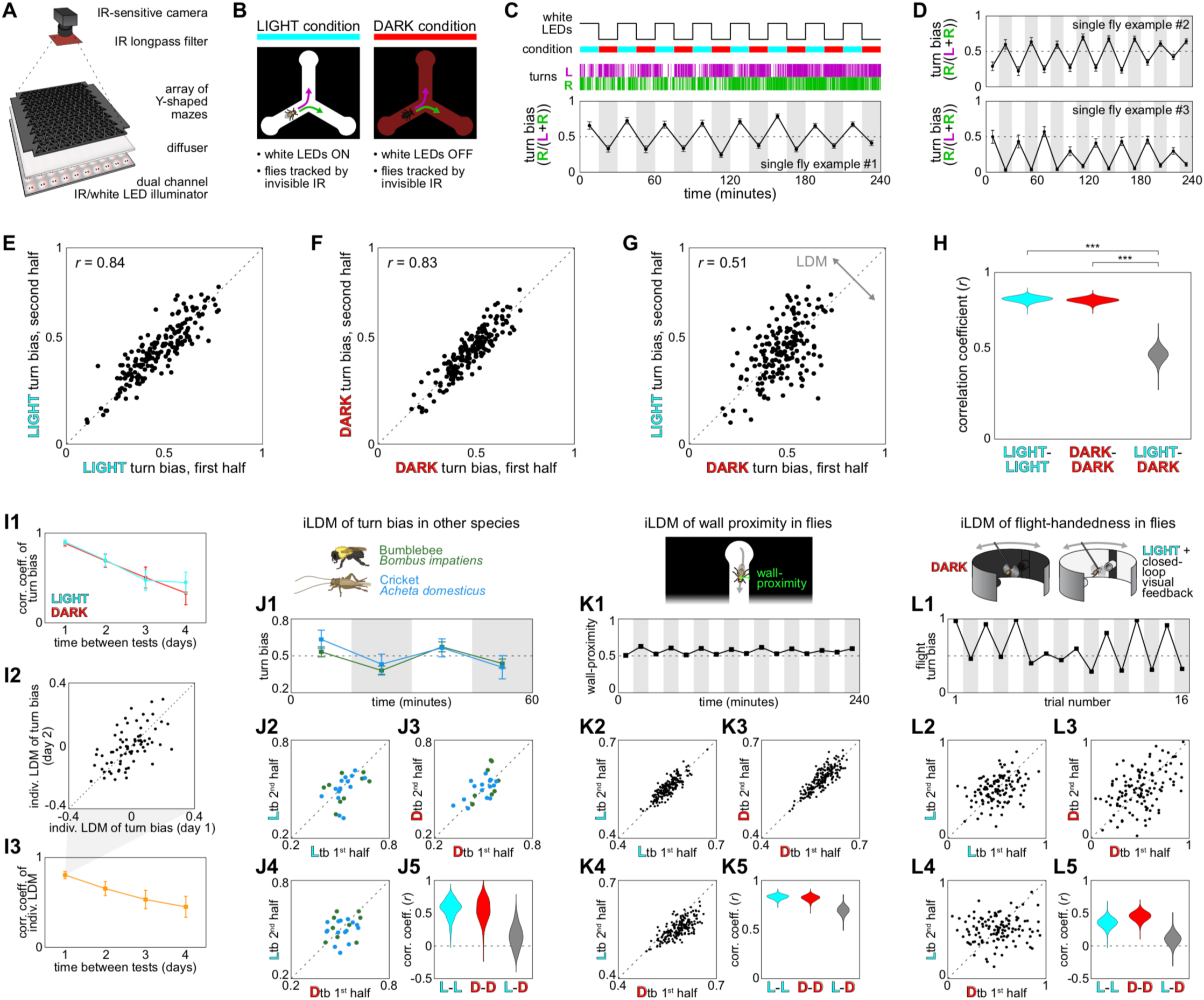
Light-dependent modulation of individual locomotor behaviors. **A**) Schematic of instrument for imaging individual flies locomoting in symmetrical Y-shaped mazes. **B**) Behavior is recorded in two luminance conditions. In LIGHT experimental blocks, white LEDs are on; in DARK blocks white LEDs are off. In both conditions, flies are tracked by constant infrared illumination. **C**) Example behavioral data from an individual fly exhibiting a large light-dependent right-to-left modulation (LDM) of its turn bias (fraction of turns through the Y-maze to the right). Magenta ticks indicate turns to the left, green right (this color scheme is used in all Figs). Bars are +/-SEM of turn bias in each block. **D**) Behavior from two other flies showing other patterns of light-dependent modulation of turn bias (left-to-right and slightly-to-strongly-left). **E**) Scatter plot of individual turn biases from LIGHT blocks in the last two hours of the experiment versus the first two hours (*n* = 197). **F**) For the same flies as E, scatter plot of individual turn biases from DARK blocks in the last two hours of the experiment versus the first two hours. **G**) For the same flies as E and F, scatter plot of individual turn biases from LIGHT blocks in the last two hours of the experiment versus DARK blocks from the first two hours. The spread of these points off the x=y line shows LDM of turn bias. **H**) Bootstrap-derived confidence distributions of the *r* values in E-G. Here and throughout the paper * = *p*<0.05, ** = *p*<0.01, *** = *p*<0.001. **I**) Stability of LDM over days: **I1**) Correlation of light and dark turn biases as a function of the number of days between tests. C.f. Buchanan et al., 2015. **I2**) Scatter plot of LDM (difference of light and dark turn biases) on day 2 vs day 1. Points are individual flies. **I3**) Correlation of LDM as a function of the number of days between tests. **J**) LDM in bumblebees (dark green) and crickets (blue): **J1**) Turn biases of individual insects across luminance conditions (comparable to C and D). **J2**-**J4**) Scatter plots of turn bias in light and dark conditions. Points are individual insects (comparable to E-G). **J5**) Bootstrap-derived confidence distributions of the *r* values in J2-4. **K**) As in J, but for LDM of wall proximity in flies. **L**) As in J and K, but for LDM of turn bias of flies mounted in a flight simulator in the dark and stimulated by at dark bar on a bright background whose position is under closed-loop control of the fly.

We exposed flies to 15 minute blocks of alternating dark and light luminance conditions over an experiment lasting four hours. We observed some flies to have strikingly different turn biases (proportion of right turns out of total turns at the choice point of the Y-maze) between the dark and light conditions (Fig 1C, D). Some flies were left-biased in the light and right-biased in the dark, or vice versa. Other flies were slightly left-biased in the light and strongly left-biased in the dark. The bias shown in each of the light condition blocks was generally consistent over the course of the experiment, as was the distinct bias shown in the dark. Across flies, the biases exhibited in the light blocks of the first half of the experiment were strongly correlated with the biases in the light blocks of the second half of the experiment (*r*=0.84; Fig 1E), and likewise for biases in the dark (*r*=0.83; Fig 1F). As expected from the examples of individual flies whose bias changed with the luminance condition, the correlation between biases in the light in the first half of the experiment and the dark in the second half was significantly lower (*r*=0.51; *p*<0.001; Fig 1G, H). The overall variances of turn bias in light and dark conditions were similar (σ^2^=0.0112 and σ^2^=0.0112 respectively; Fig S1A). We refer to the change in turn bias that individual flies exhibit between the dark and light conditions as “individual light-<u>d</u>ependent <u>m</u>odulation (LDM) of turn bias” (we will use “LDM” to refer to “LDM of turn bias” by default). To quantify the magnitude of LDM observed across a population, we calculated the standard deviation of individual LDM with a correction for the number of turns each fly made (flies making fewer turns have more sampling error in the estimation of their biases, and yield inflated LDM values without this correction).

Individual measures of fly behavior (Ayroles et al., 2015; Bierbach et al., 2017; Buchanan et al., 2015; Freund et al., 2013; Kain et al., 2012; Linneweber et al., 2019; Schuett et al., 2011; Todd et al., 2017; Werkhoven et al., 2019a) are typically stable over days. This was indeed the case for the specific turn biases that flies showed in the dark and light conditions, as well as their individual LDM values (Fig 1I). In other words, LDM is itself a persistent, idiosyncratic behavioral characteristic. Consistently, the distribution of individual LDM values was highly overdispersed compared to null models in which LDM is not idiosyncratic (i.e., all flies have identical “true” LDMs and apparent variation in LDM comes from sampling error; *p*<<10^-5 by Wilcoxon rank sum test; Fig S1B). We observed LDM over the entire range of light intensities possible with our pulse-width-modulation of LED brightness (Fig S1C). Consistent with this high sensitivity, painting over the compound eyes and ocelli with opaque black paint did not significantly diminish LDM (Fig S1D,E). Aligning turn events by the transition between luminance conditions allowed us to estimate the rate at which LDM occurs. We found that immediately after dark-to-light transitions flies showed a turn bias that was indistinguishable from their overall light bias (Fig S1F). The same rapid transition in biases was seen across light-to-dark transitions (Fig S1G). This implies that LDM occurs very quickly, though estimating precise time constants of modulation is difficult because turns happen over an extended period of time, limiting the resolution of this analysis.

To assess the generality of LDM-type effects, we measured turn bias of bumblebees and crickets in larger versions of the Y-mazes. We found that these insects also had idiosyncratic turn biases that were modulated by luminance (Fig 1J). Moreover, other individual behavioral measures were also modulated by luminance, such as flies’ average distance to the nearest wall of the Y-mazes (Fig 1K; S1H-J). Such effects on the components of walking behavior were not universal, as flies did not exhibit LDM of turn rate, a measure of average speed (Fig S1K), though turning on the light did bring about a non-idiosyncratic decrease in mean turn rate. Importantly, LDM did not appear to be limited to walking behaviors, as tethered flies flying either in the dark or in the presence of a closed-loop visual stimulus exhibited modulation of their flight handedness, as measured by the area under the rightward portion of each fly’s turn distribution (Fig 1L), suggesting that the circuits mediating LDM may be upstream of divisions between walking and flight control circuits.

### Neural circuit elements mediating LDM

Since turn bias was modulated by changes in luminance, we examined whether flies with mutations in opsin genes still exhibited LDM of turn bias (Fig 2A). *norpA* flies, which harbor loss-of-function mutations in a phospholipase-C necessary for vision, exhibited no LDM. This suggests that the LDM effect is indeed due to light rather than any experimental confound like changes in temperature or electric fields. Flies with mutations in *Rhodopsin1* (mediating luminance- and motion-vision) also showed essentially no LDM (though this effect was allele-dependent). Flies with the *eyes absent* mutation (that impairs compound eye development) and *GMR-hid* flies (that express a cell-death protein in eye cells) both exhibited large reductions in LDM, though they still exhibited some. We found no reduction in LDM in flies with mutations in *cryptochrome* or *Rhodopshin4*, an opsin mediating color vision.

**Figure 2.**
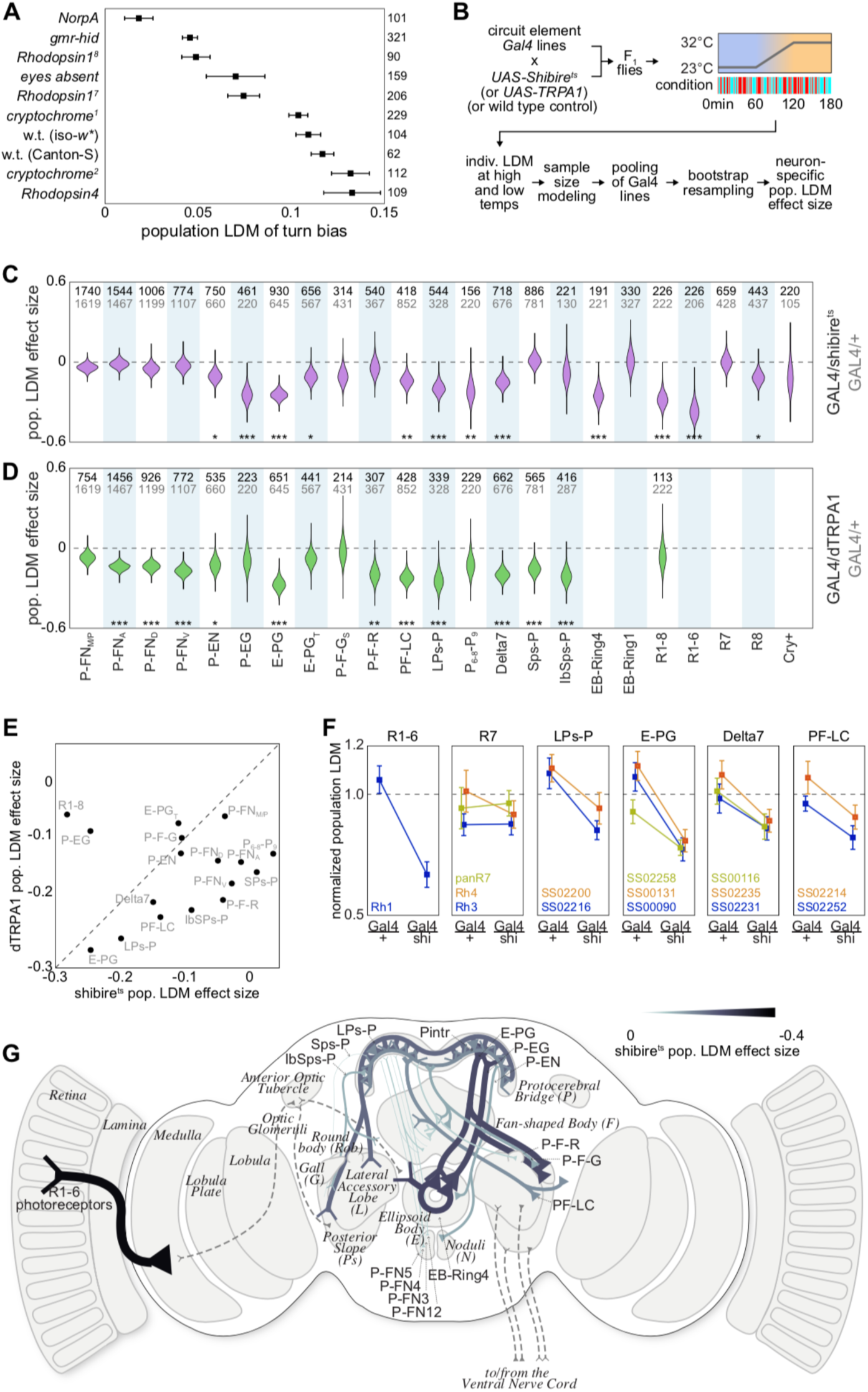
Neural circuit elements mediating light-dependent modulation of individual turn bias. **A**) Population LDM of flies harboring mutations affecting vision. LDM is significantly reduced in *NorpA, gmr-hid, Rh1*^*8*^, *eyes-absent*, and *Rh1*^*7*^ compared to wild type (w.t.) flies. Bars are +/-SE of the estimate of LDM determined by bootstrapping; numbers are sample sizes. **B**) Schematic of the thermogenetic protocol for silencing or activating specific neural circuit elements and assessing the effect on LDM. **C**) Confidence distributions of the estimated effect size on LDM from inhibiting neurons expressing Shibire^ts^ in the cell-types labeled in C, as determined by bootstrapping. 0 indicates no effect. Black numbers are the sample size of experimental animals, grey numbers control animals. **D**) As in C, but for activation of targeted neurons due to the expression of dTRPA1. Estimates in C and D aggregate multiple Gal4 lines per neuron type. See F and Fig S2. **E**) Scatter plot of median effect sizes on LDM of activating neurons with dTRPA1 vs. silencing them with Shibire^ts^. **F**) Differences in LDM between experimental animals and corresponding Gal4/+ control animals, by neuron targeted (panels) and specific Gal4 transgene (lines). Bars are +/-SE of the estimate of LDM determined by bootstrapping. Population LDM values for each experimental group are normalized by their corresponding Gal4/+ control. **G**) Diagram of the position and connectivity of the neurons targeted in this screen in the fly brain. Neuron thickness and color indicate the effect size of silencing that neuron with Shibire^ts^. Y-shaped arrow heads are postsynaptic regions, filled triangles are presynaptic. Dashed lines indicate known connections not screened. Italics labels are neuropils, roman labels are neuron types.

To identify central brain circuit elements required for LDM, we conducted a screen of transgenic lines targeting specific classes of neurons. We focused on the central complex, as recurrent connections between the protocerebral bridge and ellipsoid body neuropils within that structure harbor a representation of flies’ orientation in azimuthal coordinates (Kakaria and de Bivort, 2017; Seelig and Jayaraman, 2013) that could underpin turning behavior. These structures receive visual inputs (Omoto et al., 2017; Seelig and Jayaraman, 2013; Shiozaki and Kazama, 2017; Sun et al., 2017), as well as other sensory inputs (Pacheco et al., 2019), potentially including those bearing self-motion information (Namiki and Kanzaki, 2016; Shiozaki and Kazama, 2017), which is present in both the dark and the light, unlike visual information. We used a collection of split-Gal4 lines (Wolff and Rubin, 2018; Wolff et al., 2015) to target specific cell types covering nearly all the cell types in the protocerebral bridge (plus a selection of other Gal4 lines expressed in the ellipsoid body and the visual sensory periphery). In these cells we expressed effectors to manipulate neural activity and assess the effect on LDM. To name these cells, we follow the convention of Wolff et al., 2018: namely, a single letter for each CX neuropil to which the cell projects (multiple letters with mixed upper and lower case when there are multiple neuropil with the same first letter) separated by hyphens to indicate putative post- and pre-synaptic regions of the neuron as annotated by confocal microscopy (e.g., E-PG refers to neurons with post-synaptic sites in the EB and presynaptic sites in the protocerebral bridge and gall; Table S1). As effectors, we used the thermogenetic activator UAS-dTrpA1 (Hamada et al., 2008) or inhibitor UAS-Shibire^ts^ (Kitamoto, 2001).

F1 animals bearing Gal4 driver and effector transgenes were loaded into mazes at the permissive temperature. The luminance condition was switched between light and dark across blocks lasting between 5 seconds and 60 seconds, in a randomized order that was identical for all screen experiments. We recorded fly behavior over three temperature blocks: one hour at permissive temperature (23°C), one hour at restrictive temperature (32/29°C for Shibire^ts^ and dTrpA1, respectively), and one hour of temperature ramp between them (Fig 2B). On average, flies completed 168 and 446 turns in the permissive and restrictive phases, respectively (Fig S2A), with the mean number of turns in each phase across Gal4 lines ranging between 105 to 259 and 141 to 546. Importantly, we did not find that our thermogenetic manipulations substantially reduced activity level with the exception of one (activation of P-F-Rs through SS02293-Gal4). As seen previously (Akhund-Zade et al., 2019; Ayroles et al., 2015; Buchanan et al., 2015), the mean turn bias in all experimental groups was approximately 0.5 (0.49 +/-0.011). The variability in turn bias ranged from 0.09 to 0.15 (standard deviation) across all experimental groups (Fig S2B). The effect size of the thermogenetic manipulations on LDM was computed as the difference in the population LDM at the restrictive temperatures between experimental animals (expressing both a Gal4 and an effector) and their matched genetic controls (expressing only the Gal4), divided by the population LDM of their genetic controls. For almost every cell type, if we had multiple Gal4 lines targeting that cell type, the effect sizes were similar across the Gal4 lines (Fig 2F, S2C,D; an exception to this trend was P-E-Ns). Therefore, LDM effect sizes by cell type (Fig 2B) were calculated by pooling experimental groups targeting the same cell type.

We identified a number of central complex cell types required for LDM (Fig 2C). Using Shibire^ts^ to block vesicular release in neurons mediating the ring attractor between the protocerebral bridge (P) and ellipsoid body (E) reduced LDM by 10.67 to 24.46% (*p*=0.005 to *p*<10^−5^ by bootstrap resampling test). These neurons include the P-ENs, P-EGs, E-PGs, and Delta7s (Wolff et al., 2015). Blocking neurons carrying visual signals into the central complex, both at the sensory periphery (R1-6 photoreceptors) and proximate to the central complex (the EB-Ring4 neurons), reduced LDM by 28.29 and 24.91% respectively (both *p*<10^−5^). At least one class of input neurons to the protocerebral bridge from the lateral accessory lobe (L) and posterior slope (Ps) are also apparently required for LDM, as blocking the LPs-P neurons reduced LDM by 20% (*p*<10^−5^). Blocking the central complex output neurons, P-F-G_S_ cells, P-F-Rs and PF-LCs, also reduced LDM (10.5, 5.9 and 14% respectively; *p*=0.038, 0.16, 0.001). These output neurons connect the CX to neuropils outside that structure, including the lateral accessory lobe (L), crepine (C) and gall (G). E-PGs and P-EGs also project out of the CX and reduce LDM when silenced, as mentioned above. Notably, no perturbations of CX neurons with Shibire^ts^ increased LDM, though not all reduced it. Silencing any of the classes PFNs, which are columnar neurons within the central complex, had no effect on LDM.

We also screened most of the same neurons using dTRPA1 to assess the effects of activating these circuit elements on LDM (Fig 2D). Many of the same neurons that reduced LDM when silenced also reduced LDM when ectopically activated, including PENs, E-PGs, Delta7s, LPs-Ps, and PF-LCs. Indeed, there seemed to be a positive correlation between the effect on LDM of silencing a neuronal type with Shibire^ts^ and activating it with dTRPA1 (Fig 2E). This is consistent with previous observations that both silencing and activating neurons that control the variability of a behavior across individuals can have the same effect (Buchanan et al., 2015; Honegger et al., 2019). The most notable exceptions to this correlation were the R1-6 photoreceptors and P-EG, which strongly reduced LDM when silenced but had little to no effect when activated. Taken together, the experiments in these screens reveal a pathway of circuit elements mediating LDM of turn bias (Fig 2G) that originates at the sensory periphery (R1-6), includes input neurons to the central complex (LPs-P and EB-Ring4), the core EB-PB ring attractor subcircuit (E-PG, P-EG, P-EN and Delta7), and output neurons of the central complex (E-PG, P-EG, P-F-G and PF-LC) projecting to the gall and lateral accessory lobe, a neuropil that contains descending neurons.

### Decomposition of LDM into sensory context components

A signature of LDM is the lower correlation of individual turn biases between sensory conditions versus within sensory conditions (Fig 1E-H). Circuit elements mediating LDM show a diminished reduction in the correlation between light and dark conditions following their perturbation with thermogenetic reagents (this yields a positive LDM effect size; Fig 2). We realized this effect could happen in a combination of ways when we examined the turn bias trajectories of individual flies across the thermogenetic experiment. In some of the lines we tested, the distribution of turn biases in the light differed at the restrictive temperature, while the distribution of turn biases in the dark was roughly constant across the temperature conditions. Thus, the effect on turn bias of perturbing a circuit element could be light-specific, or dark-specific, or a combination of both (Fig 3A).

**Figure 3.**
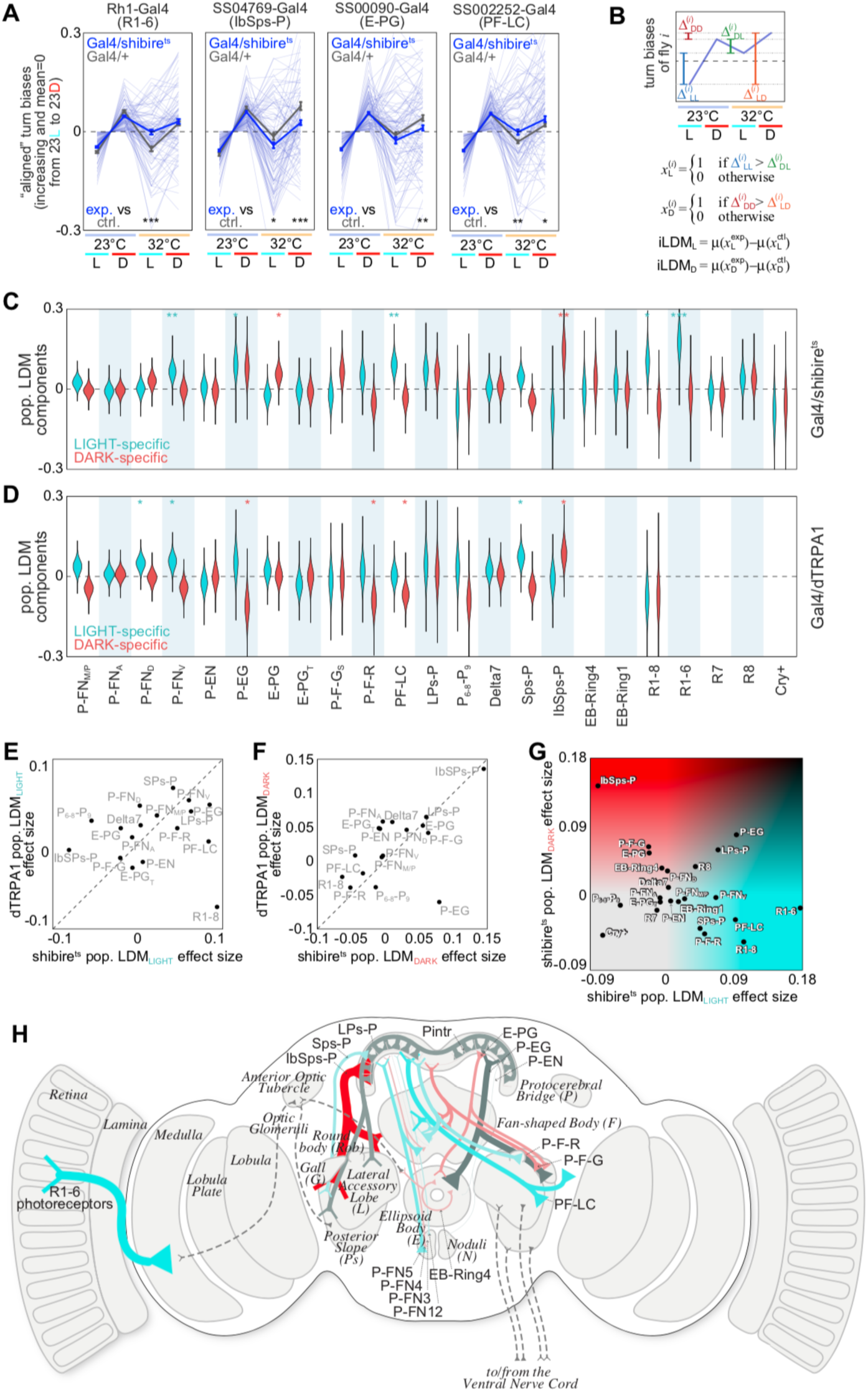
Neural circuit elements mediating sensory context-specific components of LDM. **A**) “Aligned” turn bias of individual flies (light blue) as a function of experimental block (light and temperature conditions), for representative Gal4 experiments. L=LIGHT, D=DARK. Alignment consists of subtracting the mean turn bias across the first two blocks and flipping the values of some flies so that all trajectories increase between the light to dark at low temperature. Implemented with the formula (TB_i_-mean(TB_23,L_,TB_23,D_))*(TB_23,D_-TB_23,L_/abs(TB_23,D_-TB_23,L_)). Thick grey lines represent the mean aligned trajectories of control Gal4/+ flies, thick blue lines represent Gal4/Shibire^ts^ experimental flies. Bars are +/-SEM. Differences between the control and experimental means in the light and/or dark at high temperatures indicate sensory context-specific effects of perturbing those neurons on LDM. **B**) Schematic and formulas for calculating light- and dark-specific components of LDM. *i* indexes over flies in an experimental group, µ is the mean, *x* is an individual turn bias in the (L)ight or (D)ark, Δ indicates absolute differences in turn bias between pairs of experimental blocks. **C**) Confidence distributions of the estimated light- and dark-specific components of the LDM effect of inhibiting neurons expressing Shibire^ts^ in the cell-types labeled in D, as determined by bootstrapping. **D**) As in C, but for activation of targeted neurons due to the expression of dTRPA1. **E**) Scatter plot of median effect sizes on the light-specific component of LDM of activating neurons with dTRPA1 vs. silencing them with Shibire^ts^. **F**) As in E, but for the dark-specific component of LDM. **G**) Scatter plot of neurons in the space of their dark-vs. light-specific components of the effect on LDM from silencing them with Shibire^ts^. Cyan indicates neurons whose effect is predominantly light-specific, red predominantly dark-specific, dark grey a combination of both. **H**) Diagram of the neurons screened, in the style of 2G. Neuron thickness indicates the effect size of silencing that neuron with Shibire^ts^. Neuron color indicates its dark- and light-specific components, i.e., its position in G.

Intuitively, we observed that the effect of perturbing photoreceptors was light-specific (Fig 3A). Silencing central complex neurons at times had light-specific effects and at times dark-specific effects. We developed a metric to quantify the light- and dark-specific effects of perturbing a circuit element (Fig 3B). For each circuit element, the value of the light-specific (or dark-specific) component of this metric is large when the perturbation specifically changed the turn bias exhibited in the light (or dark). We computed these two components for all the circuit elements in our thermogenetic screen (Fig 3C,D). As before (Fig 2E), either activating or silencing neurons generally revealed the same light- or dark-specific effects on turn bias (Fig 3E,F). The most conspicuous exception, in the case of light-specific effects, were the photoreceptors. Silencing them made light turn biases at the restrictive temperature more like dark turn biases at the permissive temperature. But activating them had the opposite effect; light turn biases at the restrictive temperature were more different from dark biases at the permissive temperature, as compared to control flies. Thus, activating photoreceptors with dTRPA1 effectively makes light-specific turn biases more intense.

Having identified the light- and dark-specific effects of each circuit element, we could place these neurons in a two-dimensional phase space of their effect on individual turn bias and its sensory-modulation (Fig 3G,H). At the stages of CX input pathways, local neurons, and output pathways, both sensory context-specific and context-independent neurons are evident. At the visual periphery, the photoreceptors mediate light-specific turn biases. Proximal to the central complex, Sps-P provides light-specific inputs, IbSps-P dark-specific inputs, and LPS-P context-independent inputs. The local Delta7 protocerebral bridge neurons mediate both components of bias. Among output neurons, PF-LC (and likely P-F-R) neurons provide light-specific outputs, E-PGs (and likely P-F-Gs) dark-specific outputs, and P-EGs context-independent outputs.

### Functional responses of P-F-R and PF-LC in behaving flies

Because we observed that perturbing PB output neurons (P-F-G, E-PG, P-F-R, PF-LC) decreased the amount of LDM we observed, we hypothesized that the activity of those cells may be modulated by light (especially P-F-R and PF-LCs, which showed a light-bias specific effect). However, there are multiple ways in which these neurons could affect locomotor biases. For example, they could be functionally symmetrical, but upstream of asymmetrical neurons, so that when they are activated, they contribute to motor asymmetry indirectly. Or, they themselves could be functionally asymmetrical and directly cause context-dependent behavioral asymmetries. These two scenarios make different predictions. If the behavioral asymmetries are imparted downstream of these CX output neurons then we may see consistent responses across individual animals. However, if these neurons are themselves the source of context-dependent bias, then we might observe functional asymmetries that correlated to individual LDM.

To test this hypothesis, we imaged neural activity in CX output neurons with two-photon microscopy while flies walked on an air-supported ball (Fig 4A). As in our y-maze experiments, we provided periods of time with or without ambient illumination (Fig 4B). We found that individual flies displayed locomotor handedness on the ball (Fig 4C) and that this handedness was modulated by the light, consistent with an LDM effect (Fig 4D, E). We recorded Ca^2+^ dynamics using a pre-synaptically localized variant of GCaMP6s (Fig 4F; Cohn et al., 2015). Simultaneously, we recorded ball kinematics using the FicTrac software package (Moore et al., 2014). For P-F-Rs, we imaged axonal arbors in the round body (Fig 4F, S3A). We observed variable spontaneous activity throughout the experiment in P-F-Rs. Sometimes, changes in Ca^2+^ coincided with changes in ball motion (Fig 4F). We did not observe significant changes in the mean Ca^2+^ activity during illumination (Fig 4G, H). We found a modest correlation between turning during the lights on period and the change in Ca^2+^ activity when lights were turned on (r=-0.23; Fig 4I). We did not see any correlation with forward velocity (r=0.04; Fig 4J). We saw a similar pattern with LDM of turning (Fig 4K) and forward velocity (Fig 4L). While the correlation between P-F-R responses to light and LDM of turning is not statistically significant, it is potentially consistent with the observation that silencing P-F-Rs diminishes LDM in the Y-mazes by 5.9% (Fig 2).

**Figure 4.**
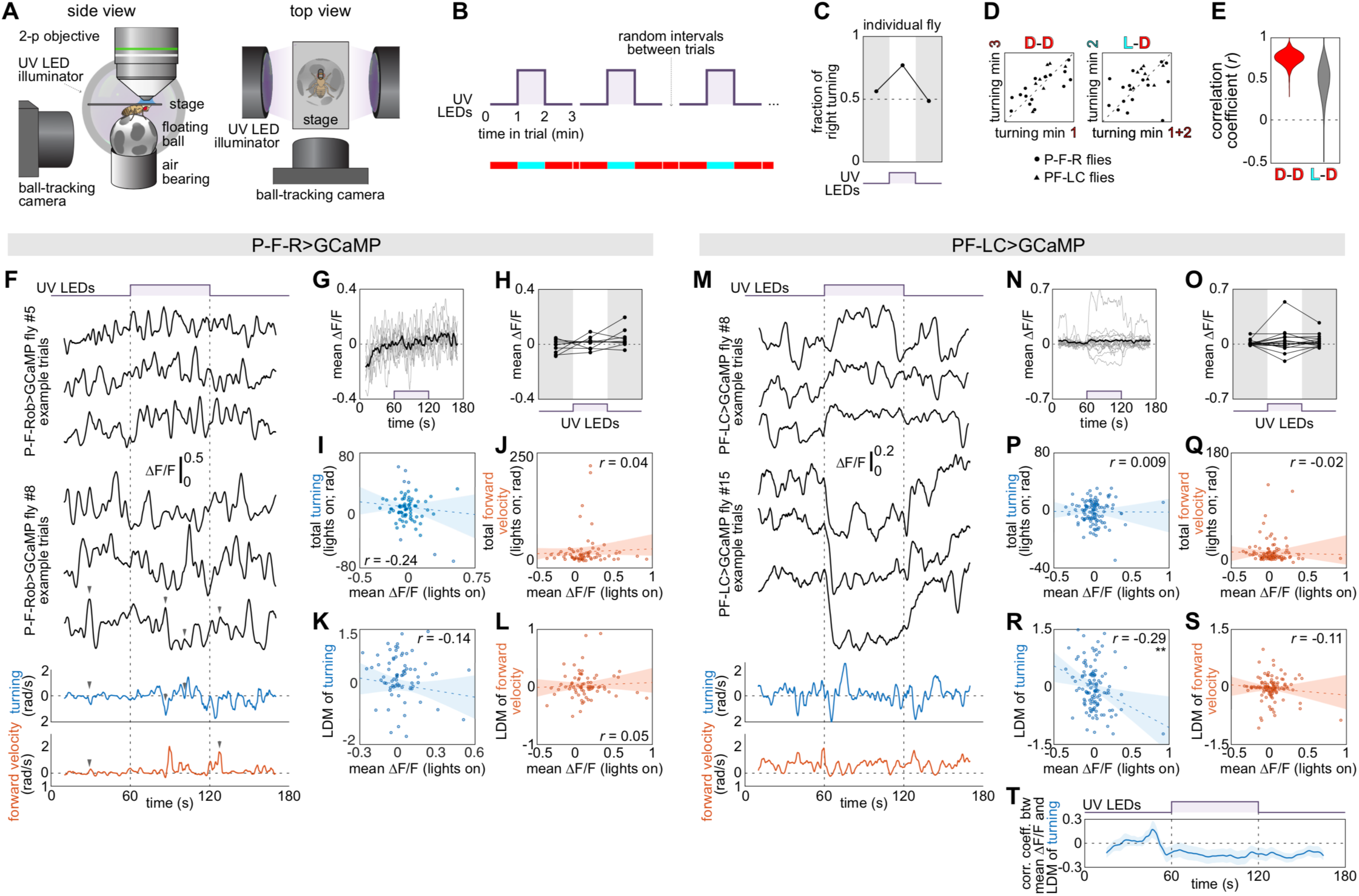
Physiological responses of central complex output neurons in behaving flies. **A**) Diagram of a fly mounted for 2-photon Ca^2+^ imaging on a floating ball, with flanking UV illuminators to change the luminance condition. **B**) Structure of the imaging trial. **C**) Turn bias on the ball, for a single fly, over the three blocks of the experiment. **D**) Scatter plots of the turn bias on the ball between the two dark blocks (left) and the lights on block vs. the average of the two dark blocks (right). Points are individual flies; shape indicates which neurons are expressing sytGCaMP6s (P-F-Rs or PF-LCs). **E**) Bootstrap-derived confidence distributions of the *r* values in D. **F**) Three example Ca^2+^ responses from the axonal bundle in the Round Body of P-F-Rs from two individual flies (See also Fig S3). Blue and red traces at bottom show reconstructed turning and forward velocity of the fly from the trial with the preceding Ca^2+^ recording. Arrowheads indicate synchronized Ca^2+^ and ball rotation spikes. Ca^2+^ traces smoothed with 2.5 second sliding average for visualization. **G**) Mean ΔF/F vs. time for individual flies (grey) and the mean of all fly means (black). *n*=9 flies. **H**) Mean ΔF/F vs. trial block. Connected sets of points are flies. **I**) Total turning during the lights on block vs. the mean ΔF/F in that block. For flies imaged in the right hemisphere, turning was flipped in sign to maintain ipsi/contra-lateral relationship. Dotted line is a linear fit; shaded region the 95% CI of that fit. Points are trials. *n*=72 trials across 9 flies. **J-L**) As in I, but for forward velocity, the LDM of turning and the LDM of forward velocity, respectively. **M-S**) As in F-L, but for Ca^2+^ responses recorded from the dorsal Lateral Accessory Lobe axonal projections of PF-LCs, in the left hemisphere. *n*=117 trials across 15 flies. **T**) Correlation coefficient between the LDM of turning and mean ΔF/F in a 5 second sliding window over time. By comparison, panel R examines the correlation of LDM of turning and mean ΔF/F over the entire lights-on block. Line is the mean correlation; shaded area is +/-SE as estimated by bootstrapping.

For PF-LCs, we imaged their axonal arbors in the dorso-lateral region of the LAL (the region with the greatest axonal innervation; Fig S3B). We observed variable spontaneous activity throughout the experiment in PF-LCs (Fig 4M); We also often observed strong individual modulation of neuronal Ca^2+^ when the lights were turned on (Fig 4N,O). For some animals, the GCaMP6s signal went up substantially across trials, for some animals down. On average across flies there was no change. Unlike P-F-R, we did not find correlation between turning or forward velocity and Ca^2+^ activity (Fig 4P,Q). However, we observed a significant correlation between LDM of turning (but not forward velocity) and change in Ca^2+^ during illumination (Fig 4R,S, *r*=-0.29). This relationship was likely not due to individual flies having different mean LDMs of turning or Ca^2+^ response (Fig S4). We found that the correlation between between LDM of turning and PF-LC activity was absent or reversed prior to the lights coming on (Fig 4T). After the lights were turned off, Ca^2+^ activity in PF-LCs continued to be negatively correlated with LDM of turning, and this may reflect the baseline shift of Ca2+ induced by the light persisting after the lights go off (Fig 4M, S3B). The correlation of Ca^2+^ activity in PF-LCs with LDM of turning suggests that idiosyncratic differences in the activity of these neurons could impart context-dependent biases onto locomotor behavior.

### Neurons post-synaptic to P-F-R and PF-LC

Although we found calcium correlates of turning LDM in PF-LCs innervating the lateral LAL, how these cells, and the LAL in general, is still largely a matter of speculation. Nine bilateral pairs of descending neurons have been identified projecting from the LAL to the VNC that could directly influence motor output (Namiki et al., 2018). However, DNs project from at least 34 other distinct brain regions in *Drosophila*. To refine our hypotheses of how P-F-Rs and PF-LCs and could be mediating a light-specific effect on locomotor bias, we sought to identify the neurons postsynaptic to these cells. To do this, we used the down-stream neuron labeling system *trans-*Tango which labels pre-(under UAS control) and postsynaptic neurons with two different fluorescent reporters (Talay et al., 2017; see Table S1).

We found that P-F-Rs made pre-synaptic connections to a large number of neurons within the central complex (Fig 5A). Especially obvious were projections in the PB, FB, and No (Fig 5B). P-F-Rs have both spine-like and bouton-like processes in the FB (Wolff et al., 2015), so we were not surprised to see *trans-*Tango staining in the FB. The labeling pattern and cell body location of many of the downstream neurons we found were consistent with P-FN morphology. These neurons have been hypothesized to store spatial memory within the central complex (Stone et al., 2017). We expected to see significant post-synaptic target cells throughout the LAL as P-F-R project to the Round Body, at the dorsal margin of the LAL. However, we saw sparse labeling in the LAL, and only in the dorso-lateral region. In addition, we saw staining in the crepine. Both of these features are diagnostic descriptors of the PF-LC class, which suggested to us that P-FRs are upstream of PF-LCs. According to anatomical descriptions (Wolff et al., 2015), P-F-Rs and PF-LCs project to adjacent layers of the FB. Thus, a direct P-F-R-to-PF-LC connection may exist at the boundary between FB layers 3 (P-F-R) and 2 (PF-LC).

**Figure 5.**
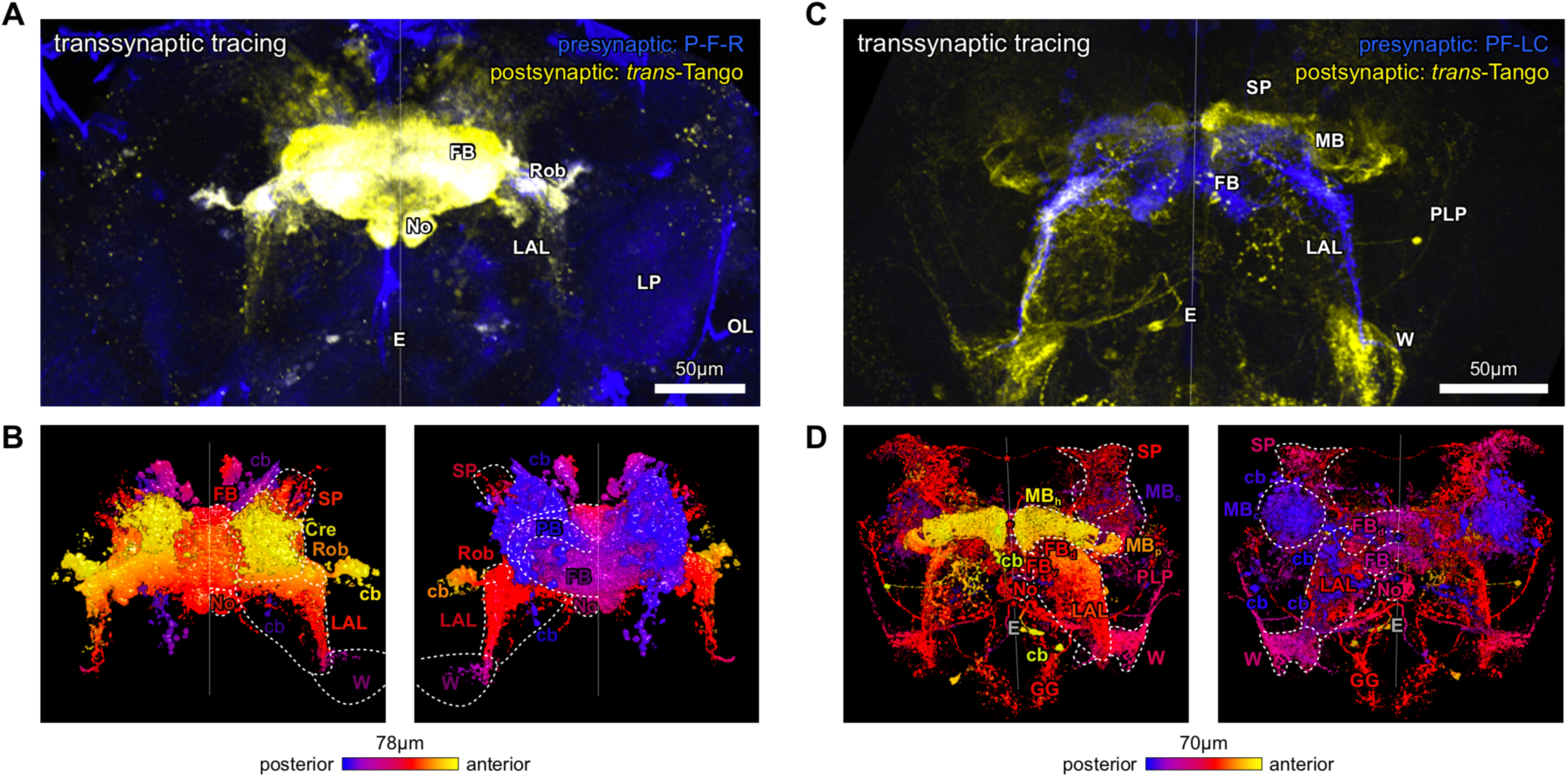
Characterization of neurons postsynaptic to P-F-R and PF-LC. **A**) *trans*-Tango labeling of neurons (yellow) postsynaptic to P-F-R (blue). Dotted line is the midline. Neuropil labels (only shown on the right): E=esophagus, FB=fan-shaped body, LAL=lateral accessory lobe, LP=lateral protocerebrum, No=noduli, OL=optic lobe, Rob=round body. **B**) Surface reconstruction of postsynaptic neurons. View from the anterior at left, posterior at right. Color indicates depth along the anterior-posterior axis. Dashed lines demarcate neuropils. Cre=crepine, W=wedge, cb=cell bodies. Label colors match the depth of the structure they label. **C** and **D**) As in A and B, but for PF-LC and its postsynaptic partners. FB_d_=dorsal FB, FB_v_=ventral FB, GG=gnathal ganglion, MB_c_=mushroom body calyx, MB_h_=horizontal lobes of the MB, MB_p_=MB peduncle, PLP=posterior lateral protocerebrum, SP=superior protocerebrum. Asterisks mark midline-crossing projections. Portions of the yellow channel of C in the gnathal ganglion were masked out of the reconstruction and cropped out of the panel because they were present in *trans*-Tango/+ control animals (Fig S5; Methods).

The neurons postsynaptic to PF-LCs (Fig 5C,D) appeared to predominantly project to neuropils other than the CX. We observed a large number of cross-hemispheric LAL neurons, which appear to match several cell types described in Franconville et al., 2018 as functionally connected to PF-LC. According to those authors, these midline-crossing neurons project to the contralateral medial LAL. In *Drosophila*, the ventromedial LAL is thought to be more involved in motor control (Namiki and Kanzaki, 2016), so these cross-hemispheric connections may have a motor role. We observed neurons projecting to the gamma lobes of the mushroom body and to the superior protocerebrum. Missing from this set of postsynaptic cells are any descending neurons or projections into the neighboring PS, another site of descending projections. However, we did find prominent labeling in the wedge, which also contains significant projections to the VNC (Namiki et al., 2018). We observed *trans*-Tango staining in the gnathal ganglia and antennal mechanosensory and motor center; however, similar staining was seen in *trans*-Tango/+ negative controls (Fig S4), so we cannot, with confidence, determine if there are neurons in those areas postsynaptic to PF-LC. From the *trans*-Tango analysis overall, we conclude that P-F-Rs are likely presynaptic to PF-LCs, PF-LCs project to many other brain regions, and neither of these cell types appears to be directly presynaptic to descending neurons.

### Correlation between PF-LC anatomical asymmetry and behavior

Because the turning biases we observed were stable over days (Fig 1I), we hypothesized that individual asymmetries in the structural anatomy of PF-LCs, like the extent of its axonal projections, could impart context-dependent motor asymmetry. Moreover, if information only leaves the CX through P-F-Rs via PF-LCs, which is possible given their apparent hierarchical relationship (Fig 5), then PF-LCs are that much more of a bottleneck, and the possibility that they are a locus of behavioral individuality may be increased. We hypothesized that asymmetries in PF-LC anatomy could asymmetrically excite downstream cells leading to motor asymmetries. The LAL, one of two neuropils where PF-LCs project, is thought to be segmented into sensory and motor areas along the dorsal-ventral axis (Namiki and Kanzaki, 2016), and PF-LCs are distributed broadly along the lateral edge of the LAL (Wolff et al., 2015; Fig 6B). Thus, we suspected that PF-LC asymmetries in certain regions of the LAL may be more predictive of locomotor bias than others. To assess the relationship between presynaptic asymmetry in PFLCs and behavior, we combined the LDM assay with synapse-localized labeling and immunohistochemistry (Fig 6C). We expressed a truncated version of the Bruchpilot (Brp) protein fused to the fluorescent reporter mStraw (Mosca and Luo, 2014) in PFLCs using SS02252-Gal4 (Fig 6B). Prior to immunohistochemistry, we recorded behavior for individual animals in Y-mazes in our behavioral instruments and measured their light and dark turn biases. Next, we dissected out their brains, stained them with antibodies targeting Brp, and collected image stacks of the pre-synaptic neuron volumes. To compute asymmetry as a function of position, we projected the Brp labeling intensity (a proxy for synapse size/count) onto the long axis (roughly dorso-ventral) of the axon bundles and then compared each bundle with its contralateral pair (Fig 6C,D; Movies S1 & S2).

**Figure 6.**
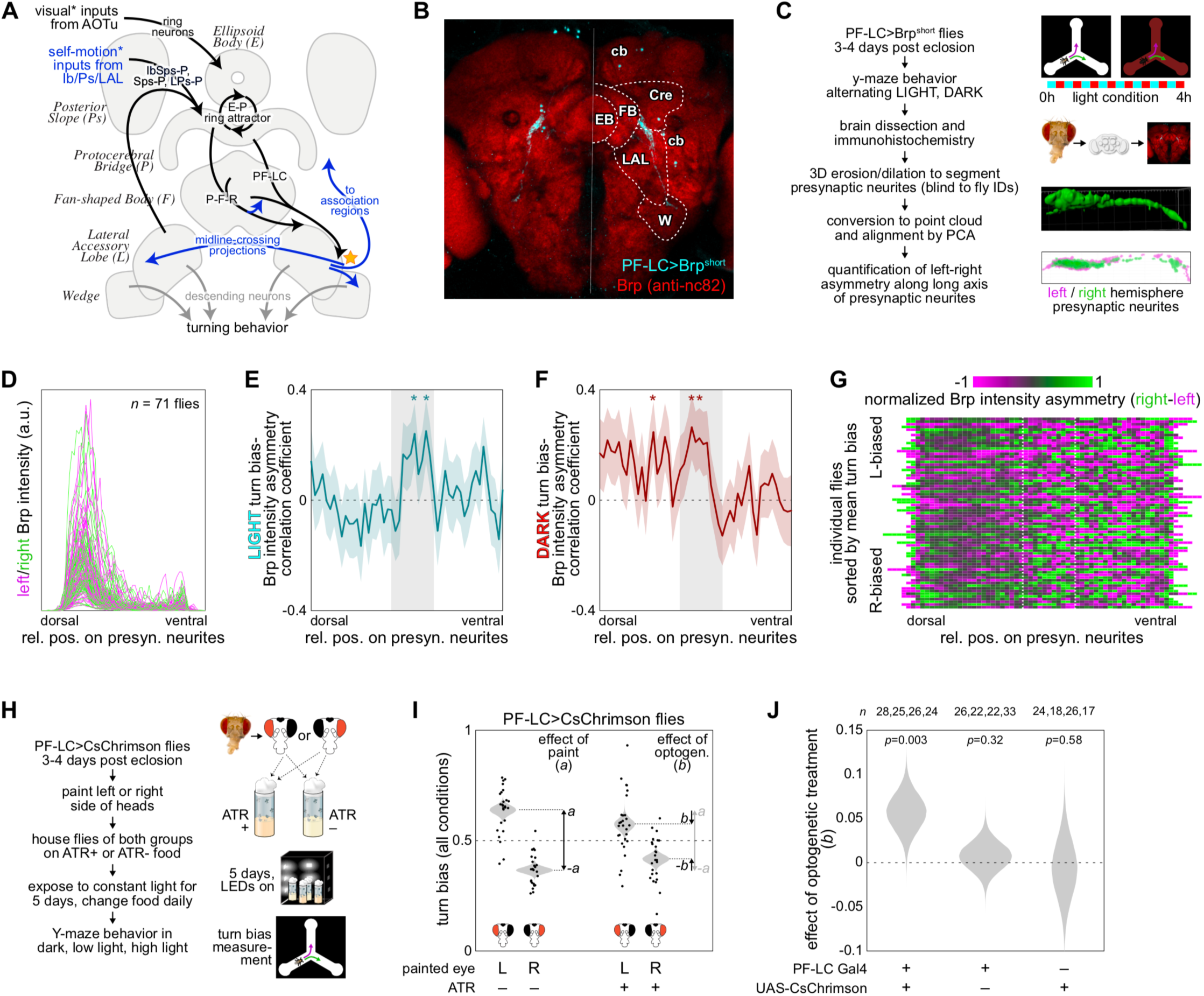
PF-LC asymmetry determines individual behavior. **A**) Diagram of neural circuit mediating light-dependent modulation of turn bias. Black elements indicate previously known pathways and functions, blue elements indicate new inferences of this work, grey elements indicate hypotheses. Orange star indicates the axons of PF-LCs as a site of bottlenecked outputs of the central complex, and potential site where individual physiological or anatomical variation could impart individual behavioral biases. * acknowledges that self-motion information likely arrives at the CX in part through the EB (Shiozaki and Kazama, 2017) and visual information in part through the LAL (Namiki and Kanzaki, 2016; Omoto et al., 2017). **B**) Immunohistochemical labeling of the presynaptic compartment of PF-LCs using the GFP-tagged Brp^short^ marker that localizes to synaptic active zones (cyan). Red is pan-neuronal Brp (*α*-nc82 antibody) as a background stain. Dotted line indicates the midline; dashed lines outline neuropils. Cre=crepine, EB=ellipsoid body, FB=fan-shaped body, LAL=lateral accessory lobe, W=wedge, cb=cell bodies. **C**) Protocol for correlating individual turning biases and left-right anatomical asymmetry in PF-LC LAL presynaptic compartments. **D**) Intensity of Brp^short^ synaptic active zone staining along the long axes of individual PF-LC LAL presynaptic compartments. Magenta is left hemisphere, green is right. **E**) Correlation between left-right asymmetry of Brp^short^ intensity in PF-LC LAL presynaptic compartments and turn bias exhibited in the light, as a function of position along the long axis of the projections. Cyan shaded region is +/-SE of the correlation as determined by bootstrapping. Grey shaded region demarcates a zone of higher correlation also present in F and marked in G. **F**) As in E, but for the correlation between PF-LC LAL asymmetry and the turn bias exhibited in the dark. **G**) Left-right asymmetry of individual PF-LC LAL pairs as a function of position, sorted by ascending turn bias (average of light and dark biases). White dotted lines demarcate the grey shaded region of E and F. **H**) Protocol designed to induce left-right asymmetries in PF-LC using chronic optogenetic stimulation. **I**) Turn bias of PF-LC>CsChrimson flies painted on the left or right eye (and ocelli) and stored in constant illumination for 5 days. Flies subject to this treatment and not fed ATR showed altered turn biases (away from the side of painting; *a*). Flies fed ATR exhibited an additional change in turn bias (*b*) in the opposite direction, presumably due to the optogenetic stimulation. Grey densities indicate the confidence around the estimate of the mean turn bias for each group. Points are individual flies. **J**) Bootstrap-derived confidence distributions for the effect of optogenetic stimulation of PF-LCs on turn bias (*b*). Numbers indicate the sample sizes of the 4 experimental groups shown in panel I that were used to calculate each distribution. These effects were seen at multiple luminance levels during behavioral testing of experimental animals, but not genetic controls (Fig S7).

We calculated the correlation coefficient between individual turn bias in the light or dark and Brp intensity asymmetry along the long axes of PF-LC axonal bundles (Fig 6E-G). We found a significant positive correlation near the middle of the axonal bundle with both light and dark locomotor biases (*r* ≅ 0.2 and 0.25, respectively). This positive relationship means that greater PF-LC presynaptic volume in the right hemisphere is associated with more right turning (Fig 6G). As an alternative measure of neuronal asymmetry, we quantified left-right differences in only the total volume of the Brp-positive voxels. This measure had a similar spatial profile of correlation with turn bias, but was less positive (Fig S6A,B). By this measure, there was a second region of positive correlation between PF-LC axonal asymmetry and turn bias in the light, closer to the distal tip of the axonal bundle, perhaps indicating that PF-LC asymmetry in the wedge is predictive of an individual’s light bias. We also noticed that asymmetry of Brp staining density (intensity divided by volume) was positively correlated over the whole axonal arbor with the turn bias exhibited in the dark (Fig S6C,D), but the biophysical significance of this measure is not immediately obvious.

### Inducing asymmetry in PF-LCs causes asymmetry in behavior

It was possible that the correlations between PF-LC activity and turning LDM, and PF-LC anatomical asymmetry and turn bias are not causal. To rule out this possibility, we devised an experiment to induce asymmetry in PF-LCs experimentally. If their relation to locomotor asymmetries were causal, inducing a left (or right) mean asymmetry in a group of animals should induce a corresponding lateralized change to the mean turn bias. We subjected flies to chronic, asymmetric optogenetic activation of PF-LCs by painting half the head of flies expressing UAS-CsChrimson in PF-LCs. We hypothesized that constant activation of these cells over days would induce physiological effects that would persist after optogenetic stimulation was removed, and that these effects would be asymmetric because the paint would shade one hemisphere more than the other. We housed flies prepared in this manner in constant white LED illumination for five days before recording locomotor biases (Fig 6H). In control flies (not fed the CsChrimson cofactor all-trans-retinal, ATR), we observed a change in the mean turn bias in the direction of their unpainted side (µ=0.14 when the left side was painted; Fig 6I). This effect was apparently not phototactic since we observed it in flies behaving in the dark (Fig S7). In the experimental flies (fed ATR),we also observed a mean shift in the direction of the unpainted side, however, the magnitude of that shift was decreased (µ=0.08; Fig 6I). This decrease was significant compared to genetic controls not expressing CsChrimson (*p*=0.003; Fig 6J, S7), and we interpret it as an optogenetic-specific effect of size 0.06 in the opposite direction of the effect of painting alone (Fig 6I). These results demonstrate that asymmetrically perturbing PF-LCs causes a mean asymmetry in turn bias. This complements our early finding that symmetrically perturbing PF-LCs with thermogenetics causes an on-average symmetric change in the variance of turn bias (Figs 2,3).

## Discussion

That individual animals behave idiosyncratically is a familiar observation to scientists studying behavior (and often a frustration when trying to measure an experimental effect). This variability represents real biological signal but is understudied and the biological mechanisms underlying individual variation (particularly within a population of animals with matched genetics and environment) are largely uncharacterized. We investigated the origins of sensory context-dependent individuality in a simple locomotor behavior. Specifically, we found that individual flies display different turn biases (predispositions to turn left or right in a Y-shaped maze) depending on the presence or absence of visible ambient illumination. This light-dependent modulation altered several (but not all) dimensions of locomotor behavior, in walking and flying flies, and in other insect species (Fig 1), suggesting it is a general phenomenon.

We found that neurons in visual pathways and the central complex mediate this modulation (Fig 2), and that their roles could be decomposed to light-specific and dark-specific effects (Fig 3). CX output neurons included both light-bias mediating and dark-bias mediating neurons, and we recorded Ca^2+^ activity in two populations of light-bias mediating output neurons, P-F-R and PF-LC, while flies walked on a ball and were subject to changes in illumination. PF-LCs exhibited idiosyncratic offsets in Ca^2+^ when the light came on, and this correlated with their individual light-dependent modulation of turning on the ball (Fig 4). Consistently, we observed that individual left-right anatomical asymmetry in the axonal arbors of PF-LC predicted individual turn bias, and inducing functional asymmetries in PF-LCs using chronic optogenetic stimulation induced behavioral asymmetry (Fig 6).

Handed locomotor biases at the individual level have been found to be stable over time in numerous species including humans (Lebovich et al., 2019) and flies (Ayroles et al., 2015; Buchanan et al., 2015; Linneweber et al., 2019). The stability of these biases over time suggests a rigidity to individual behavioral biases. But here we found that individual flies have different biases in the light and dark (Fig 1C), consistent with the idea that innate behaviors can still be “cognitively” flexible (Gorostiza, 2018). LDM is not specific to walking, as we found flies in tethered flight also displayed LDM (Fig 1L). It is also not specific to fruit flies, as it was observed in both crickets and bumblebees (Fig 1J). Goal-directed walking behaviors likely rely on internal representations of heading (Corfas et al., 2019; Green et al., 2019; Kim and Dickinson, 2017) that are updated by external visual cues (Turner-Evans and Jayaraman, 2016). To maintain an accurate estimate of heading, insects are thought to combine self-motion information with visual cues (Shiozaki and Kazama, 2017; Sun et al., 2018). It is possible that LDM arises from discrepant biases associated with these two sensory streams, with one bias displayed in the dark when the animals rely on self-motion information and a different bias displayed in the light when the animals use both self-motion and visual information. A simple prediction of this hypothesis is that depriving the animals of one of these sensory streams will reduce the change in bias accompanying the change in luminance. We observed that chronic or induced disruption of the visual system, specifically Rh1-expressing photoreceptors and the *norpA* mutation, decreased LDM significantly. Painting the eyes of the fly with opaque black paint or using very dim light, unexpectedly, did not reduce LDM (Fig S1E). Therefore, we do not think that LDM depends on image-formation. Instead, illumination without visual detail may be sufficient to induce LDM, but we cannot rule out the possibility that our eye-painting failed to cover tiny, but sufficient, portions of the image-transmitting eye.

Neurons of the CX, notably the E-PG cells, have been shown to encode heading, an important idiothetic feature for path integration, while walking (Green et al., 2017; Seelig and Jayaraman, 2015; Turner-Evans et al., 2017) and in flight (Giraldo et al., 2018) and appear to contribute to ring attractor dynamics (Su 2017, Kakaria 2017). The position of the bump of activity in EPGs that encodes heading can be updated with or without visual feedback, and we hypothesized they could be involved in LDM. In general, we found that both silencing or activating neurons in the PB decreased LDM (Fig 2C-G). Previously, we investigated an *in silico* version of the heading system and found that both increasing or decreasing activity of several classes of neurons disrupted normal ring attractor function in the circuit (Kakaria and de Bivort, 2017). Silencing or activating PF-LCs decreased LDM to a similar degree as E-PG neurons. It is striking that every significant effect of a thermogenetic manipulation reduced LDM. We suspect that this is due to the construction of the LDM metric, which derives from the variance of individual context-dependent changes in turn bias. If LDM ultimately arises from something like the sum of independent stochastic effects distributed across circuit elements that are differentially activated by light, then the variances of these effects will sum positively, and disrupting any of them will reduce the total variance. Which circuit elements contribute more variance to LDM likely depends on their intrinsic predispositions to vary stochastically and functional positions in the CX circuit.

While heading is encoded in the PB and EB, the behavioral features represented in other CX neuropils is a matter of ongoing inquiry. Due to the columnar projections from the PB, the FB likely inherits heading signals in some form from the PB, so both post-synaptic compartments of PF-LCs and P-F-Rs (the PB and FB) likely encode a bump. However, the pre-synaptic axons of PF-LCs in the LAL are likely intermixed (Turner-Evans and Jayaraman, 2016), and there is no conspicuous spatial organization of P-F-R axons within the compact Round Body. These cell types may then encode scalar values rather than a vector in azimuthal coordinates. We found that presynaptic Ca^2+^ in P-F-Rs and PF-LCs was modestly correlated with turning and LDM of turning, respectively. This means that the activity of these neurons encode signals that are at least partially in motor coordinates, and that the CX accomplishes a transformation of signals from sensory coordinates, to egocentric azimuthal coordinates, to motor coordinates across a circuit that might be as compact as three layers (e.g., Ring neurons, E-PGs, PF-LCs). The position of PF-LCs as a CX-to-pre-motor bottleneck, means that they may have an outsized influence on motor programs. Stochastic variation in these neurons may therefore be particularly likely to introduce idiosyncratic biases into behavior.

The PB receives inputs from at least two distinct sources, the EB and the posterior slope (PS; Namiki and Kanzaki, 2016; Wolff et al., 2015), and this multisensory convergence likely explains the role of PF-LC in sensory context-dependent behavioral modulation. The EB has primarily been thought of as a visual neuropil (Omoto et al., 2017; Seelig and Jayaraman, 2013; Sun et al., 2017), though it has been shown to carry self-motion information, at least during flight (Shiozaki and Kazama, 2017). The PS, on the other hand, receives a large number of ascending inputs directly from the ventral nerve cord (Hsu and Bhandawat, 2016), positioning it well to transmit proprioceptive and mechanoreceptive signals that could be integrated to produce a self-motion-based estimate of heading. We suspected that these different PB input populations might differentially influence behavior in the light and the dark. Consistent with this, we observed that the IbPs-P neurons had a specific effect on the turn bias exhibited in the dark (Fig 3C,D,H), however, silencing IbPs-P neurons did not decrease LDM substantially (this may be due to a non-significant opposing effect in the light; Fig, 2C). Neurons affecting LDM in one sensory context or the other (or occasionally both) were found throughout the central complex. Surprisingly, we found that different neurons within the PB-EB ring attractor subcircuit affected specific biases. For example, activating or silencing E-PGs affected the dark bias but not the light bias. We had thought that the PB-EB ring attractor circuit would perform the same computation on sensory inputs irrespective of their origin. This suggests that the PB-EB circuit may contain functionally insulated pathways that are sensory-modality specific.

During homing (returning to the nest after foraging) in bees, CPU1 neurons (PF-LCs in *Drosophila*) have been proposed to direct steering by computing an error between an intended heading and current heading (Stone et al., 2017). To estimate current heading, the bee integrates visual stimuli and self-motion signals. Both of these sources of heading estimates may carry transient errors or persistent, lateralized biases arising in left-right asymmetries of the circuits bearing these signals. PF-LCs are downstream anatomically (Wolff et al., 2015) and functionally (Franconville et al., 2018) of neurons that carry both heading estimates and sensory information. If the difference in activity in PF-LCs between light and dark represents the discrepancy between vision- and self-motion-derived estimates of heading (i.e., the discrepancy between the position of a visible goal cue, and the representation of the egocentric position of that goal, stored, in the dark, in the bump position) then it may also be proportional to the motor correction the animal should make when the sensory context changes, in order to maintain a constant heading in both conditions.

We observed that the activity of PF-LCs, when the light came on, was predictive of the change in turn bias that the animals exhibited when the light came on. This relationship is consistent with thermogenetic perturbation of PF-LCs specifically affecting the light-component of LDM, and the correlation between PFLC activity and turning LDM being absent before the light is turned on (Fig 4T). The light-specific baseline value of PF-LC Ca^2+^ varied from fly to fly (Fig 4M,N). This was true even as the animal turned through large sweeps of azimuthal orientation. Thus, PF-LCs in the light encode a feature that is more stable than heading, such an estimate of persistent error associated with visual inputs or the offset of the bump and a visible goal (Seelig, 2015).

Activity in P-F-Rs correlated to some extent with turning over the whole trial, as well as in moment-to-moment alignment of Ca^2+^ and motion spikes (Fig 4F,I). The relationship between P-FR Ca^2+^ responses in the light and LDM of turning was not significant, but trended in the same direction as that of PF-LCs. This lack of significance could reflect the relatively low statistical power associated with our physiology experiments compared to our behavior experiments, or the lower effect size of perturbing P-F-Rs on LDM. Either way, it seems likely, given our trans-Tango tracing of postsynaptic partners of P-F-R, that its effect on turn bias, and the sensory-modulation thereof, are mediated by downstream PF-LCs.

We did not observe any descending neurons to be directly downstream of PF-LCs (Fig 5). Instead, the connection between PF-LCs and motor circuits may be made by multiple midline crossing neurons and/or neurons projecting to the lateral accessory lobe and wedge that are postsynaptic to the PF-LCs. Also downstream of PF-LCs were neurons that appeared to project to association areas in the superior protocerebrum and the gamma lobes of the mushroom body, which is the site of visual inputs to that neuropil (Vogt et al., 2016). These connections, which appear to travel away from motor areas, could potentially transmit efference copy signals about motor or heading states back to sensory and association regions.

Based on 1) PF-LCs position at the bottleneck of central complex outputs; 2) their likely encoding of a scalar value in each hemisphere in the LAL; 3) the effect on light-specific component of bias when they are thermogenetically perturbed; and 4) their activity in the light being correlated with the LDM of turn bias, we hypothesize that PF-LCs are a site where individual functional asymmetries impart idiosyncratic bias onto individual behavior — a locus of individuality. We measured the light and dark turn biases of many flies expressing the synaptic active zone marker Brp^short^ in PF-LCs and then quantified, as a function of position along a D-V axis, the left-right asymmetry of the LAL compartment of PF-LCs. We employed this spatial approach because there is a proposal that activity in the LAL is assorted along a dorsolateral-ventromedial in *Drosophila* axis into more sensory and more motor regions (Namiki and Kanzaki, 2016). Additionally, the sign of any correlation between variations in PF-LC and behavior will depend on the net sign of the intervening connections between PF-LC and motor output. This could vary spatially across PF-LC, as different populations of postsynaptic neurons likely connect to PF-LC in different regions. We found that near the middle of the LAL compartment, left-right asymmetry was modestly predictive of turn bias in both the light and dark conditions (Fig 6E-G). PF-LCs come in at least 3 subtypes, short, medium and long (Wolff et al., 2015), defined by how far they project along the lateral margin of the LAL. Our data suggest that asymmetry in medium length PF-LCs could correlate best with turn bias in general, while asymmetry in the dorsal LAL, potentially in short PF-LCs, may be correlated with turn bias in the dark (Fig 4F).

If the correlations between PF-LCs and turn bias reflect a causal relationship, it would fit with this understanding of the circuit: when the lights come on, there is an idiosyncratic change in the offset of Ca^2+^ activity in PF-LCs that drives more or less activity through the left-right asymmetric LAL compartments of PF-LC. This presynaptic asymmetry drives an asymmetry in behavior as long as the light is on, altering the activity baseline of PF-LCs. To test if PF-LC asymmetry is causal of behavioral asymmetry, we used optogenetics to stimulate PF-LCs over five days. This stimulation was made asymmetric by shading one side of the head by painting one eye and the top of the head. We surmised that this manipulation would lead to asymmetric long-term changes in the physiology of PF-LC (e.g., through classical synaptic strengthening via LTP in the post-synapse or outgrowth of PF-LC itself via activity-dependent plasticity; Ueno et al., 2013; Budnik et al., 1990; Fernández et al., 2008). We measured the turn bias of these animals in the dark and the light, and found that in all conditions, painting the left side resulted in a leftward shift in turn bias and vice-versa (Figs 6I,J and S7). Thus, ectopic PF-LC asymmetry appears to be linked to behavioral asymmetry in both the light and dark (Figs 6E,F and S5C). This appears to be at odds with our behavioral data that indicates the effect of silencing PF-LCs is light-specific. Though we did observe that activating PF-LCs with dTRPA1 selectively altered the dark-component of LDM (Fig 3D), and if changes in baseline activity of PF-LCs are light-specific (Fig 4), presynaptic asymmetry in PF-LCs will have a light-specific effect on behavioral biases.

Our test of the causal relationship between PF-LC asymmetry and behavioral asymmetry used optogenetic stimulation to induce an asymmetry. But this does little to illuminate the specific natural fluctuations that may be leading PF-LC to function asymmetrically and drive idiosyncratic sensory-modulation of behavior. These fluctuations could be stochastic lateralized differences in synapse number (consistent with our Brp imaging), post-synaptic channel number, quantal number or content, volume of presynaptic active zones, or intrinsic physiological properties. Identifying and validating these mechanisms represents a stimulating and challenging future direction for this work, and a potential path to an integrated, multi-causal level understanding of the biological basis of individual behavioral bias, and its modulation by sensory context.

## Methods

### Lead Contact and Materials Availability

This study did not generate any unique reagents. *Drosophila* lines used in this study are available upon request. See Table S2 for a complete list of lines and their source. Further Information and requests for resources and reagents should be directed to and will be fulfilled by the Lead Contact, Benjamin de Bivort.

### Data and Code Availability

All data needed to reproduce our analyses (behavioral centroid data, imaging ROI traces and micrograph images) as well as all our analysis scripts are available at *http://lab.debivort.org/light-dependent-modulation/* and on our analysis code is hosted at: *https://github.com/kskakaria/ldm_project*. Unless otherwise noted, all experiments and analysis was performed with custom written code in MATLAB 2013a-2019a.

### Experimental Model and Subject Details

#### Drosophila melanogaster

Fly lines were housed on standard Bloomington type A cornmeal-dextrose food made in the fly media facility at the Harvard Biological Laboratories. Unless otherwise noted, flies were reared in standard conditions, maintained at 23C and ∼40% relative humidity year round with 12h/12h light/dark cycle with ambient white LED illuminators.

#### Thermogenetic flies

For behavioral experiments, all experimental flies were seeded in bottles with 25mL of lightly yeasted fly food with 10 female and 5 male parental animals. After 48 hours, we transferred parents into new bottles for 48 additional hours and then discarded them. For experimental crosses, we collected virgins from *UAS* parental lines and crossed them to *split-Gal4* or *Gal4* males. On a daily basis, we collected F1s and stored them in cohorts of 16 females and 14 males 4-6 days. For the screening experiments, crosses were kept for 24 hours in a food vial to allow all virgin females to be fertilized before successively transferring all parents to bottles. To standardize genetic backgrounds, we used *UAS-Shibire* and *UAS-dTrpA1* flies outcrossed for 10 generations into a common lab strain (Honneger, 2019). Split-Gal4 lines were already in a common genetic background from their recent construction (Wolf et al., 2018).

#### Ca^2+^ Imaging

For Ca^2+^ imaging experiments, all *split-Gal4* males were crossed to *UAS-tdTomato; UAS-sytGCaMP6s* virgins.

#### Trans-Tango

We conducted *trans*-Tango experiments using flies expressing UAS-myrGFP and QUAS-mtdTomato-3xHA on the second chromosome, and elav-hGCGR::TEVcs::QF and UAS-Glucagon on the third chromosome obtained from the Bloomington *Drosophila* Stock Center (#77124, Bloomington, IN, USA). We crossed SS02252-Gal4 (PF-LCs), SS02192-Gal4 (P-F-Rs), and Iso^KH11^ males to *trans*-Tango virgins. To reduce off-target proteolytic cleavage and QUAS-mtdTomato-3xHA expression, we reared the crosses in vials in 18°C incubators for 5 weeks (3 weeks until eclosion and 2 weeks to allow sufficient protein expression).

#### Bumblebees

Bumblebees were a gift from James Crall (originally obtained from BioBest®) and were stored in miniature nest housing and fed a diet of artificial nectar and pollen before testing in Y-mazes. Bumblebees assayed for behavior were female workers.

#### Crickets

We obtained crickets of mixed sex (*Archeta domesticus*) from PetsMart (160 Alewife Brook Pkwy, Cambridge, MA 02138) and immediately tested them in Y-mazes designed for bumbleebee experiments.

### Method Details

#### Y-maze behavioral experiments

We performed all behavioral experiments in custom acrylic Y-maze imaging instruments with Sigmacote lubed acrylic lids (to prevent flies from walking upside down; Sigma-Aldrich: SL2-25ML) similar to those described in Buchanan et al., 2015. However, we used custom manufactured dual channel white/IR LED illuminator panels to facilitate light versus dark experiments (Knema, LLC). We tracked fly centroids using custom software in MATLAB (Werkhoven et al., 2019b). Turns were scored as right or left by determining the change in y-maze arm that individual flies occupied entering and exiting the decision point. To obtain higher throughput, we collected data from 4 behavioral boxes simultaneously with independent PCs allowing us to run 480 flies at a time. We controlled LED panels for all boxes using a single Arduino Uno running custom control software interfaced with MATLAB via 4 MOSFET transistors implementing pulse-width modulation to set the overall light intensity. We synchronized the start of the experiments using MATLAB, communicating via UDP signals sent from the computer controlling the Arduino Uno. By default, we set each board to a 25% PWM duty cycle as we found that to be an effective stimulus intensity (Fig S1).

For persistence experiments, we recovered flies after initially testing them in Y-mazes like normal using light anesthetization on ice. They were stored individually in standard media vials until being retested on later days as normal and then returned to individual housing. Only flies that survived the entire 4 days are shown in Fig S1J. For cricket and bumblebee LDM experiments, we created a larger version of the standard Y-maze geometry in laser-cut acrylic. It measured 8mm high, with corridors 10mm wide and 20mm long. These animals were centroid-tracked in the normal behavior instruments using the same software.

To determine wall-proximity, we analyzed the same wild type behavioral data from Fig 1C-H. We isolated x-y centroids for an individual in the approach before making a turn (only including centroids once the animal turned around at the end of the maze and moved back towards the center of the maze). From all the centroids of an animal, we calculated a boundary that marked the extent of the fly’s path in x and y. We then interpolated these boundaries to obtain dense sampling and calculated the Euclidean distance in pixels to the closest boundary point both on the left and right wall (relative to the fly). To compensate for spatial distortion caused by the relative angle from the camera we then normalized the distance for each fly as a proportion of distance between the two walls (e.g., 1 if the fly is always on the extreme right side of the maze arm preceding a right turn). For each turn we calculated the mean ipsilateral proximity (i.e., the distance from the left wall before a left turn and right wall before a right turn). To obtain an individual fly’s overall wall proximity score we then took an average over all turns.

In the experiments to measure LDM of flight-handedness, flies were suspended in a cylindrical arena with a rigid pin tether. We measured each fly’s wing-beat angles in real time, and used these signals to rotate the arena in closed-loop. The cylinder featured a vertical black stripe on a white background, which was only visible when the arena lights were on. The arena also contained two translucent tubes facing the fly: one aligned with the vertical stripe, and one 180 deg from the stripe. In the data presented here, no air was flowed through the tubes. The arena covered all of the fly’s visual field in azimuth, and most of it in elevation. Turn rates, calculated as the difference in wingbeat angles multiplied by a constant gain, were saved at 50 Hz. To calculate handedness from this data, we first found the distribution of turn rates for each trial. We then calculated the area under the turn rate distribution for rightward turns only, and divided this by the total area under the curve. This yielded a fraction from 0 to 1 for each trial, where 0 was a trial where the fly only turned left, and 1 was a trial where the fly only turned right. We plotted turn bias as a function of trial and condition for each fly. We also found the average turn bias for the 1st and 2nd halves of light and dark trials for each fly. For additional details on the flight simulator experiments, refer to Currier & Nagel, 2018.

#### Thermal manipulations

We ran Y-maze experiments in a Harris Environmental System room that controlled temperature and humidity. Thermogenetic experiments consisted of three blocks: 60 minute permissive temperature (23°C), 60 minute ramp to restrictive temperature, and 60 minute restrictive temperature. Restrictive temperature was 29°C for dTrpA1, and 32°C for Shibire^ts^. We allowed a long ramp period to allow sufficient time for heating the relatively low thermal conductance of the acrylic mazes, as well as the room as a whole. During screen experiment we cycled the lights on and off in a pseudo-random order (randomly generated originally but repeated for every experiment) of various time periods (between 5 seconds and 60 seconds). We quantified LDM as described below. To quantify effect sizes of LDM in thermogenetic screen (Fig. 2), we first mean centered and variance normalized each of turn bias distributions, separately for light and dark and each genotype. We then subtracted LDM at the restrictive temperature of controls from experimental flies and divided by LDM of controls. To quantify light bias or dark bias effect sizes of LDM effects (Fig. 3) we made pairwise comparisons between turn bias at high temperature and low temperature. We subtracted the bias observed at the high temperature condition (light or dark) for each individual from the biases observed for both low temperature conditions (light and dark). For each condition (light or dark) we then binarized the differences for each individual (smaller difference=1, larger difference=0) and averaged across individuals to obtain a population-level proportion score (LDM_L_ or LDM_D_; Fig 3B). This is one of several methods of decomposing the light- and dark-specific components of LDM that gave similar qualitative results.

#### Chronic asymmetric activation

To paint fly heads, we anesthetized 2-3 day old females on CO_2_ and applied black paint (Carbon Black Acrylic, Golden^®^) to half of the head, aiming for a complete covering of the area from the eye to the ocelli, inclusive, along the top of the head. For optogenetic activation, we used UAS-CsChrimson, which requires <u>a</u>ll-*<u>t</u>rans*-retinal (ATR) as a cofactor. ATR was added to the food in vials by pipetting 10uL of 100mM ATR in 70% ethanol onto the media surface. We transferred 30 painted flies into each ATR+ (experiment) or ATR- (control) food vial, 15 left painted and 15 right. To provide chronic optogenetic activation, we placed vials in a light isolated box with constant illumination (∼50µW/mm^2^) at 21°C (room temperature). ATR is not stable for long in the light, so we changed the vials every day to maintain optogenetic potential. After 5 days, we loaded flies into Y-mazes and tracked them as lights cycled through 5 minute blocks of darkness, low-light, and high-light (0, 20, and 75 PWM, respectively) at 21°C. We then quantified locomotor bias for each individual, in each light block.

#### Immunohistochemistry

For antibody staining for anatomical imaging, we adapted an immunohistochemistry protocol from the Janelia Research Campus FlyLight protocol for anti-GFP staining in adult brains (*https://www.janelia.org/project-team/flylight/protocols*). Briefly, we anesthetized adult flies with CO_2_, dipped them in ethanol for 10 seconds to reduce hydrophobicity of the cuticle, and rinsed them in 1x phosphate buffered saline (PBS). We transferred flies to a Petri dish of chilled bath saline and dissected them. To fix brains, we transferred them to individually labelled 0.5mL polypropylene tubes with 50uL 2% paraformaldehyde (PFA) and allowed them to fix for 60 minutes on a nutator at room temperature. We removed PFA and added 50uL of 0.5% Triton X-100 detergent in PBS (PBT) to wash and permeabilize the tissue followed by four 15 minute washes, on a nutator at room temperature. To block non-specific highaffinity binding sites, we removed PBT and added 50 uL of 5% Goat Serum diluted in PBT to each tube for 90 minutes. We removed the block and added primary antibody solution (detailed below). We returned the brains were to the nutator at room temperature for 4 hours to allow binding, and then moved them to a nutator at 4 degrees Celsius for 48 hours. We removed primary solution, and washed brains with a four times with PBT for 15 minutes. We added secondary antibody mix and allowed binding for 4 hours at room temperature before moving brains to a nutator at 4 degrees Celsius for 72 hours. Finally, we washed brains with PBT four times for 15 minutes to remove excess antibody from the tissue. We cleared stained brains in Vectashield (H-1000, Vector Laboratories, Burlingame, CA, USA) and mounted them on frosted microscope slides (12-544-2, Fisher Scientific, Pittsburgh, PA, USA). We performed all imaging at the Harvard Center for Biological Imaging on a Zeiss LSM 700 Confocal Microscope. As the volumes were captured, the investigator performing the imaging and assessing its quality was blind to the behavioral score associated with each individual brain.

#### Calcium imaging

We performed two-photon imaging experiments using a custom-built galvo-galvo microscope (Honegger, 2019) with a stage that mounted the fly on a 2d treadmill floating ball, which was supported by house compressed air. The air flow rates were tuned before each experiment to allow an individual fly to move naturally on the ball. We supplied whole-field illumination with 405nm LEDs diffused and directed by a ground glass condenser lens (Thorlabs: ACL2520U-DG6, LED405L) to fill the majority of the visual field on the left and right sides of the fly’s head. We controlled LEDs with an Arduino Uno via MOSFET transistors, with both lights being on or off simultaneously. The 2-photon beam was generated using an ultrafast Ti:sapphire laser (Spectra-Physics Mai Tai) laser tuned to 930nm. We captured Ca^2+^ dynamics using the ScanImage 2013 software. We performed all experiments with 20mW laser intensity. We imaged most flies at 64 × 64 pixel resolution at ∼18 Hz, and a few flies (PF-LC 1-3) at 128 × 128 pixels at ∼9 Hz. Each animal was in the experiment for ∼16 trials consisting of 3 minutes of laser exposure and 2 minutes +/-a random offset (gaussian distributed with 15s standard deviation) of rest (5 minutes each, ∼80 minutes total). We recorded behavior using an IR sensitive camera placed parallel to the body axis, behind the fly. Ca^2+^ imaging and behavioral acquisition were run simultaneously in two instances of MATLAB on the same computer, synchronized using TCP/IP protocol signals sent between the two instances, and the delays for these initializations were subtracted from the final time stamps. We smoothed each image with a 2d, 1.25 pixel standard deviation Gaussian filter. Pixels were smoothed in time with a 1 second moving average. To correct for x-y motion, we registered frames onto an average reference frame of the first trial for a fly using a FFT cross correlation method (Guizar-Sicairos et al., 2008) calculated on the red channel (anatomical marker) and applied to both green and red. To select ROIs, we first fit the pixel values of the reference frame with a gaussian mixture model with 2 modes designed to separate background and neuron-associated pixels and selected the top 50% of pixels in the brighter gaussian as a mask (Fig. S4X). Fluctuations were calculated as the frame-by-frame median of the values after masking. Both channels were normalized by dividing each by their respective means. The green channel was normalized by the red channel on a frame-by-frame basis to compensate for motion. To analyze calcium responses, we obtained a ΔF/F by first subtracting and then dividing by the first 10 seconds of each trial. In Ca^2+^-motion correlation analyses, we then subtracted from each trial’s ΔF/F trajectory the mean of values in the 20s prior to lights-on.

To track behavior of the fly we floated a 6.89mm diameter white cellulose acetate ball (CAS-ALA-1.3, Cospheric) on a stream of compressed air. We tuned the air flow by hand for individual flies until they moved the ball freely. We determined the rotation of the ball offline using the FicTrac software package (Moore et al., 2014; *http://rjdmoore.net/fictrac/*). We quantified turning as the negative of ball yaw to obtain a fly-centered metric. We quantified turn bias on the ball as the total turning to the right divided by the total turning in either direction minus the total turning to the right divided by the total turning in either direction. We then calculated individual LDM as the turn bias in the light minus the dark.

#### Trans-Tango

Immunohistochemistry and confocal imaging were performed as described above and in Table S1. Tango-channel image stacks were used to reconstruct postsynaptic neurons as follows: stacks were loaded into MATLAB. Every 3 z-layers, the linear brightness and contrast adjustments needed to separate positive Tango staining from background were determined, and linear interpolations of these values were applied as brightness and contrast adjustments to intervening layers. For the PFLC reconstruction the gnathal ganglion and AMMC non-specific staining were manually masked out of the image stack. Brightness and contrast-adjusted stacks were binarized, gaussian blurred with a standard deviation of ∼10 microns, and binarized liberally again to create a mask that included all the Tango-staining but excluded stray non-specific pixels. Masked stacks were exported from MATLAB and imported into UCSF Chimera (Pettersen et al., 2004) where a depth-coded colorscale was applied and the surface rendered.

### Quantification and Statistical Analysis

#### Y-maze behavioral analysis

For all Y-maze experiments, individual locomotor bias was quantified by dividing the number of right turns by the total number of turns, as described in Buchanan et al., 2015. To estimate individual LDM, we subtracted the turn bias in the light from the turn bias in the dark. For population LDM, we first compute light and dark turn biases for all flies and then calculated:

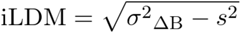

Where ΔB is the difference between the turn bias in the light and the dark and *s* is an estimate of sampling error. Subtracting an estimate of sampling error was necessary as observed LDM scales with the number of turns individuals performed. This was especially important when comparing distributions where flies perform few turns. To estimate *s*^2^, we computed the average sum of binomial variance across the light and dark:

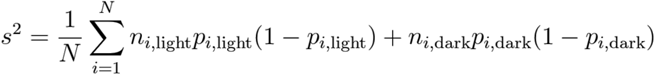

Where *N* is the number of flies, *n*_*i*,light_ is the number of turns an individual fly made in the light and *p*_*i*,light_ is the turn bias of an individual in the light (and likewise for the dark condition).

Correlation coefficients were calculated as non-parametric Spearman rank correlations. We estimated standard errors and confidence distributions using bootstrap resampling with 10,000 repetitions, with the exception of the sliding window analysis in Fig 4T, which used 100 repetitions. Reported *p*-values are nominal and have not been corrected for multiple comparisons.

#### Null modeling

To estimate null distributions of our behavioral metrics (turn bias, wall proximity and turn rate) we used resampling to estimate the standard error associated with each individual’s metrics. For turn bias, because each turn was binary (either left or right) we estimated sampling error starting with the variance of a binomial distribution (*n*\**p*\*(1-*p*)) where *n* is the number of turns and *p* is the turn bias an individual exhibited. For wall distance we estimated standard errors using bootstrapping 100 times. Individual ipsilateral wall proximities (before each turn; see above) were resampled up to the number of choices made by each fly and the overall wall proximity was calculated. The standard deviation of the distribution of wall proximities for each individual across these estimates was the estimate of individual sampling error. The standard error of turn rate for each individual was calculated by making a pool of all the inter-turn intervals from each fly, and drawing values at random from this pool while the total of its randomly sampled inter-turn intervals was less than the experimental block duration. The turn rate was then calculated for each sequence. This process was repeated 100 times and the standard deviation of the resultant distribution was the sampling error for that individual. Once we had individual sampling errors, to estimate the null distributions, we generated individual scores for each metric by sampling from a normal distribution. For each sample, the mean was the grand mean across all flies (e.g., 0.49 for turn bias) and the standard deviation was a vector of standard error values, obtained as described above, for each fly in the data set. We repeated this process 10,000 times to densely capture the distribution of possible outcomes if the distribution we observed arose only from sampling error, but flies still had the individual-specific error associated with their specific number of observations.

#### PF-LC anatomical analysis

Image stacks of PF-LCs were analyzed in 3D using Imaris 9.0 (Bitplane). First, we adjusted the minimum voxel intensity threshold to include the entire axonal projection volume. Next, we applied repetitions of dilation and erosion to denoise the 3D projection. We imported the eroded image series into MATLAB using the BFopen toolbox, and converted them into 2 vectors, point clouds (x,y,z coordinates) and intensity vectors (brightness of voxel). We aligned these images onto the long axis by mean subtracting to center on the origin and rotating them onto their principal components. Neurons that were oppositely oriented along PC1 (the long axis) were programmatically flipped, with manual inspection and correction. To compare neurons along their length we aligned the first and last 2% of each neuron’s length. The points were then divided into 50 equally spaced bins, multiplied by their intensity values and voxel volumes, summed, and smoothed with a gaussian kernel with 1 bin standard deviation. Asymmetries were then calculated by subtracting the left bins from the right bins and dividing by the sum of the left and right. To compare volume asymmetry to behavior, we then calculated a correlation coefficient for each bin with different behavioral biases.

## Supplier Index

DSHB: Developmental Studies Hybridoma Bank, Iowa City, Iowa, USA
Aves: Aves Labs, Inc., Tigard, OR, USA
TF: ThermoFisher Scientific, Eugene, OR, USA
Bio: Biorbyt, Cambridge, England
SA: Sigma Aldrich, Merck KGaA, Darmstadt, Germany
BD: BD Biosciences, San Jose, CA, USA

## Acknowledgements

We thank Ed Soucy, Brett Graham, Adam Bercu and Joel Greenwood of Harvard’s CBS Neuroengineering core for help fabricating our instruments, including our 2-photon microscope. Tanya Wolff and Gerry Rubin kindly shared the significant collection of split-Gal4 lines that we used in the circuit screen. Bryan Song and Dragana Rogulja also kindly shared Gal4 lines and mutants targeting the visual system. Tanya Wolff and Katrin Vogt provided expert consultation on the neurons in the *trans*-Tango staining. The Harvard Center for Biological Imaging, and namely Doug Richardson, were instrumental in providing resources and expert advice in all confocal imaging. Jennifer Erickson, Jess Kanwal, and Kate Leitch helped edit the manuscript. KSK and ZW were supported by NSF Graduate Research Fellowships #DGE2013170544 and #DGE1144152; TAC was supported by the NIH under grant no. R00D-C012065. BdB was supported by a Sloan Research Fellowship, a Klingenstein-Simons Fellowship Award, a Smith Family Odyssey Award, a Harvard/MIT Basic Neuroscience Grant, the NSF under grant no. IOS-1557913, and the NIH under grant no. MH119092.

## Supplementary information

**Figure S1.**
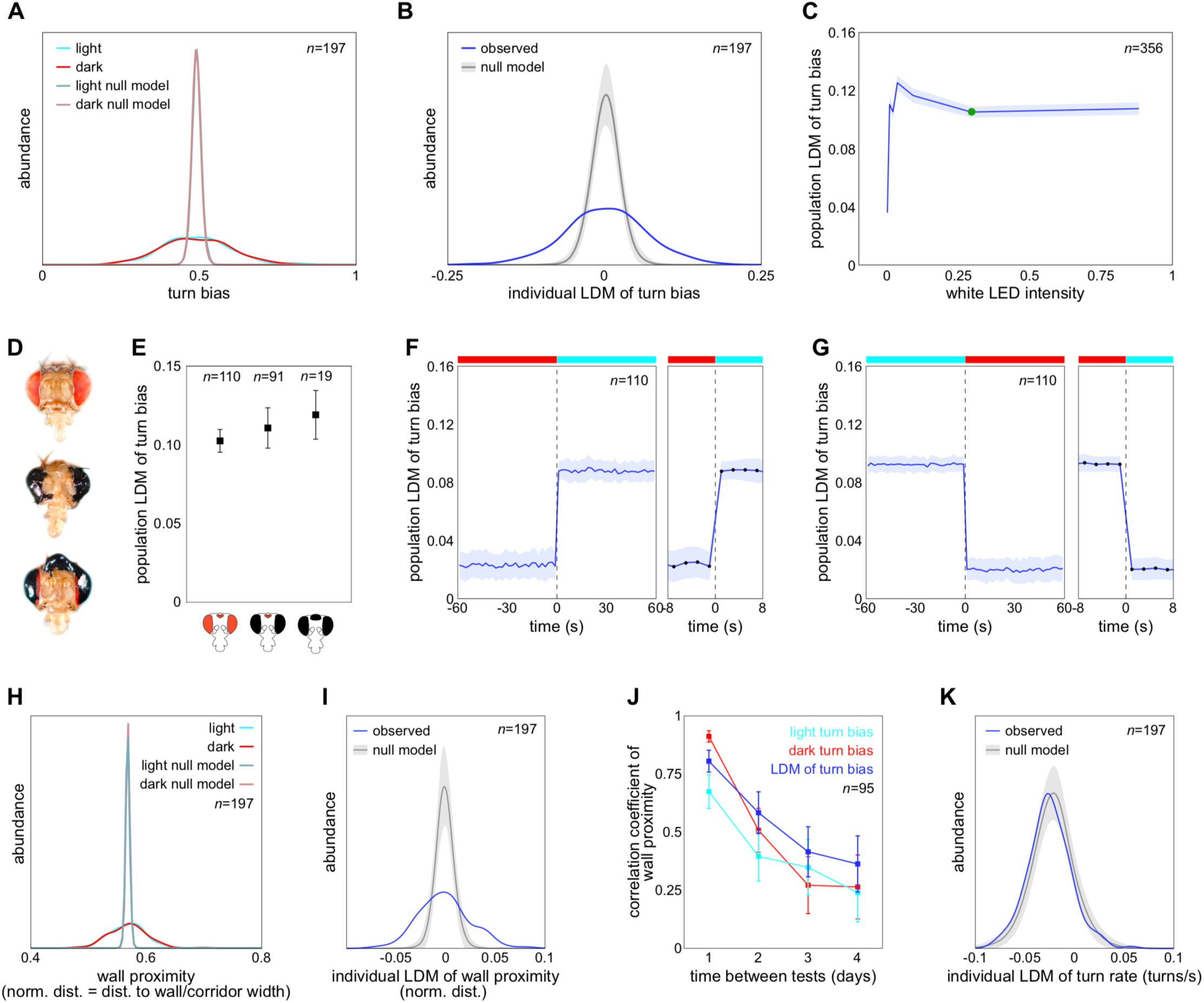
Characteristics of individual light-dependent modulation of behavior. **A**) Kernel density estimates of the distribution of turn biases in the light (cyan) and dark (red), and null models of each in which all flies choose left-vs-right with identical probabilities, and dispersion comes from sampling error alone. **B**) As in A, but for LDM of turn bias instead of turn bias, i.e., the change in individual turn bias when the light condition changes. Shaded region is +/-SE of the null model estimate, given the sample size. **C**) LDM as a function of white LED intensity (PWM duty cycle). Shaded region is the +/-SEM. Green dot indicates the conditions used throughout the rest of the paper. **D**) Photos of flies with various patterns of paint over their eyes. **E**) LDM of flies with eyes painted. Flies with both eyes and/or ocelli painted exhibit LDM comparable to unpainted flies. No paint flies plotted from Fig 1E. Bars are +/-SEM **F**) LDM as a function of time with respect to the dark-light transition. LDM is calculated over a 2 sec sliding window. Shaded region is +/-SEM. Inset at right is a zoom-in on time=0. **G**) As in F, but for the light-dark transition. **H, I**) As in A and B, but for wall proximity as a behavioral measure. **J**) Day-to-day persistence (correlation coefficient) of light bias, dark bias, and LDM of turn bias as a function of the interval between tests. Bars are +/-SE of the correlation coefficient, estimated by bootstrapping. **K**) As in B and I, but for LDM of turn rate, showing a lack of individual modulation compared to the null model.

**Figure S2.**
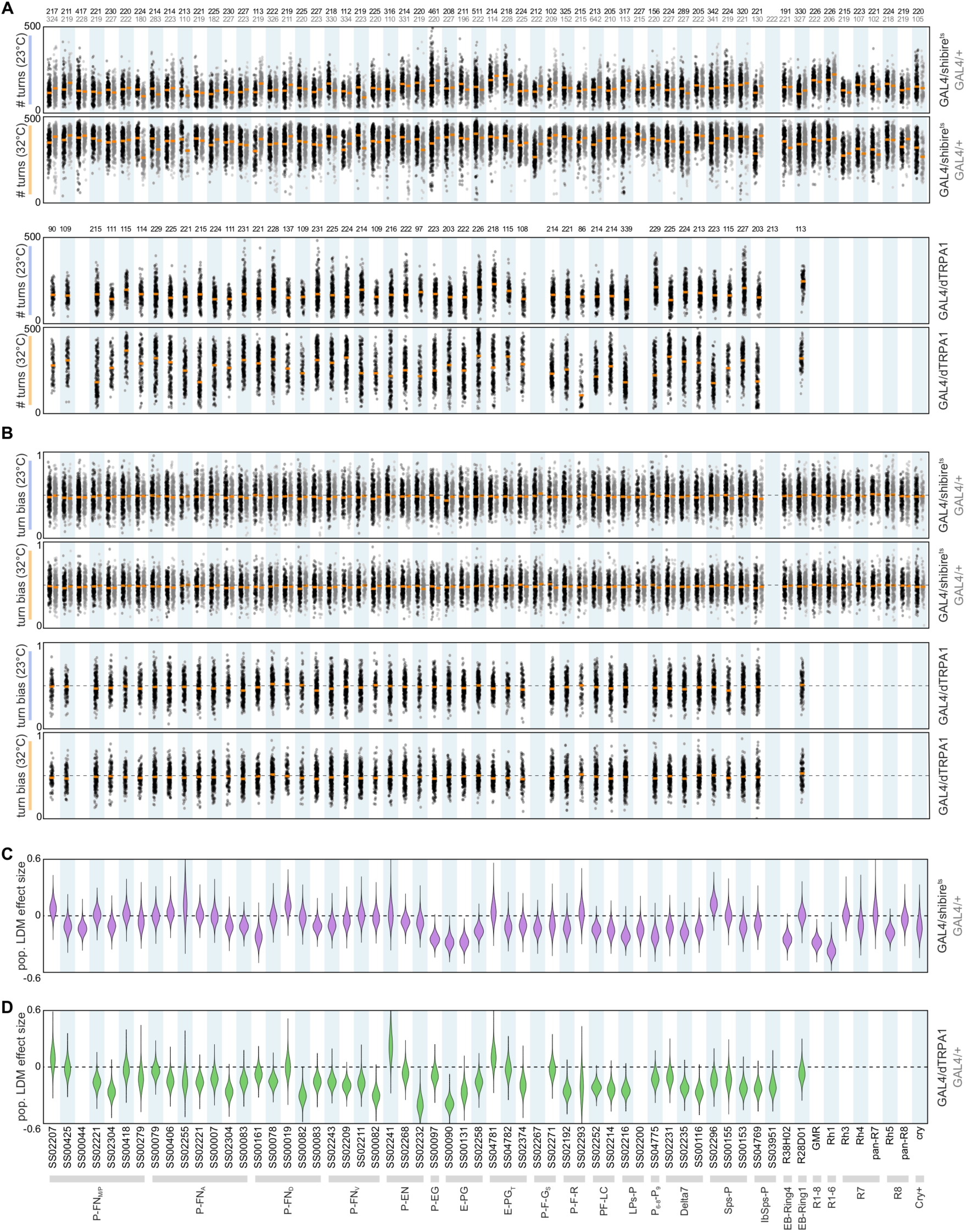
Behavioral metrics of individual flies from the thermogenetic screens. **A**) number of turns produced by individual flies (points), by Gal4 line (columns) Experimental flies in black, control flies in grey. Orange bar indicates the median of the distributions. Shibire^ts^ experimental animals top two panels, dTRPA1, bottom two. First and third panels are at the permissive temperature, second and fourth the restrictive. **B**) As in A, but for turn bias instead of number of turns. **C**) Bootstrap-derived confidence distributions for the LDM effect size of switching to the restrictive temperature for Shibire^ts^-expressing flies. **D**) As in C, but for dTRPA1-expressing flies. At the very bottom are the Gal4 names and the neuron types they label.

**Figure S3.**
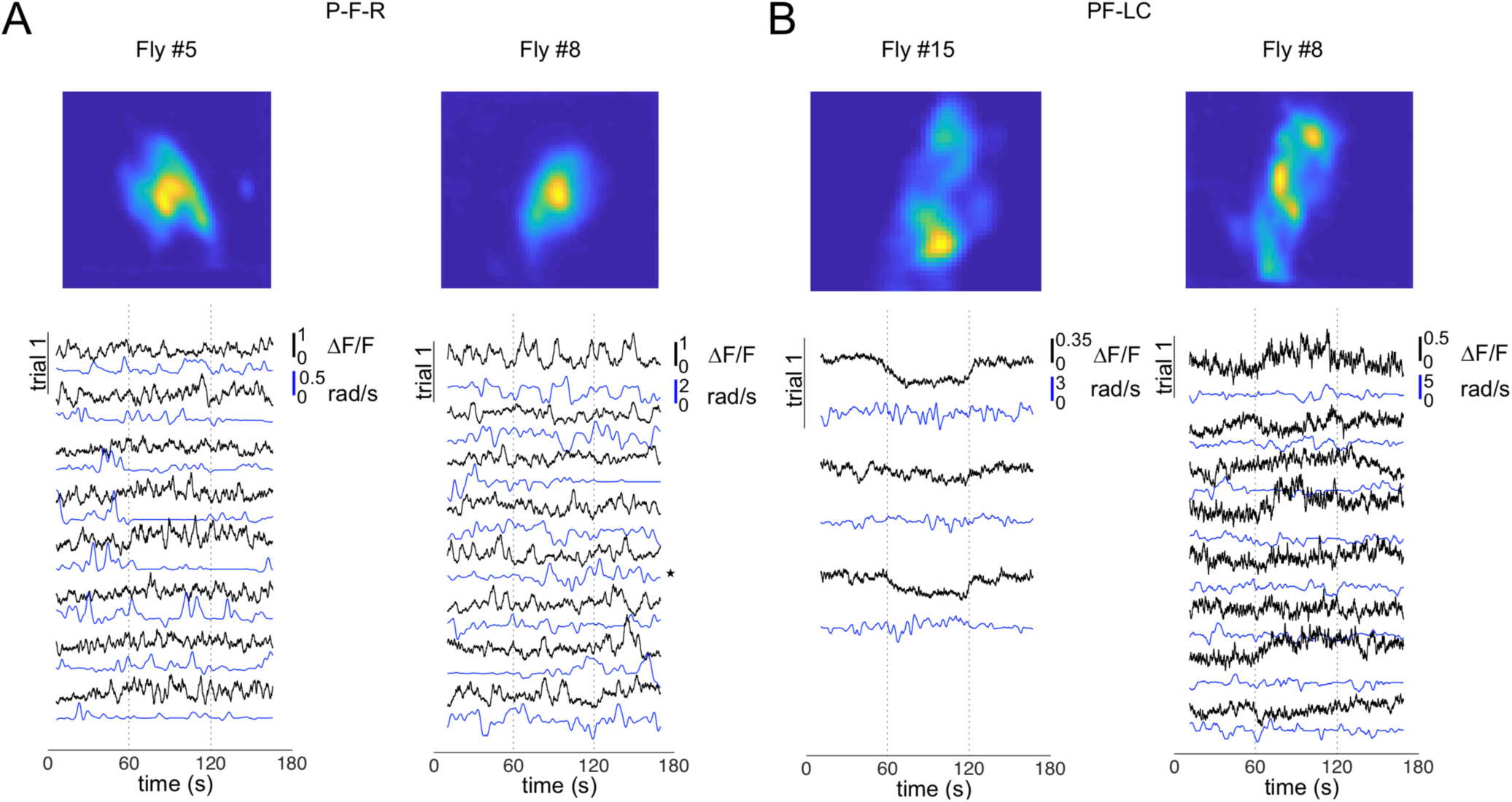
Examples of raw data from Ca^2+^ imaging experiments. **A**) The normalized mean green channel frame (syt-GCaMP6s) for P-F-Rs (SS02192-Gal4) with the imaging field of view over the Round Body, calculated across all frames, after registration (top). Raw data traces for each fly for both ΔF/F of Ca^2+^ activity (black) and ball yaw (blue). Each trial is presented as a pair of traces, with the first trial at the top. The recordings and images are from the same flies as those presented in Figure 4. Images are from the right (#5) and left (#8) hemispheres of their respective animals. **B**) Same as A but for PF-LCs (SS02252-Gal4) with the scan field of view over the dorsal portion of the Lateral Accessory Lobe. Both images are from the left hemisphere of their respective animals.

**Figure S4.**
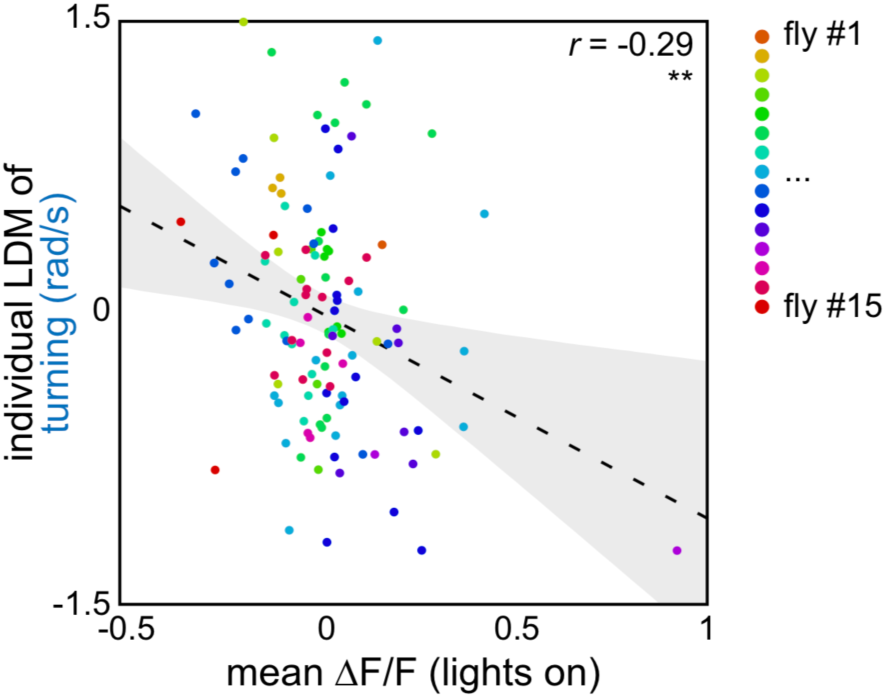
Relationship between LDM of turning and Ca^2+^ response, by fly. LDM of turning versus mean ΔF/F during the lights on block, with the trials from each individual in a distinct color. The negative correlation does not appear to arise by different mean ΔF/Fs across flies.

**Figure S5.**
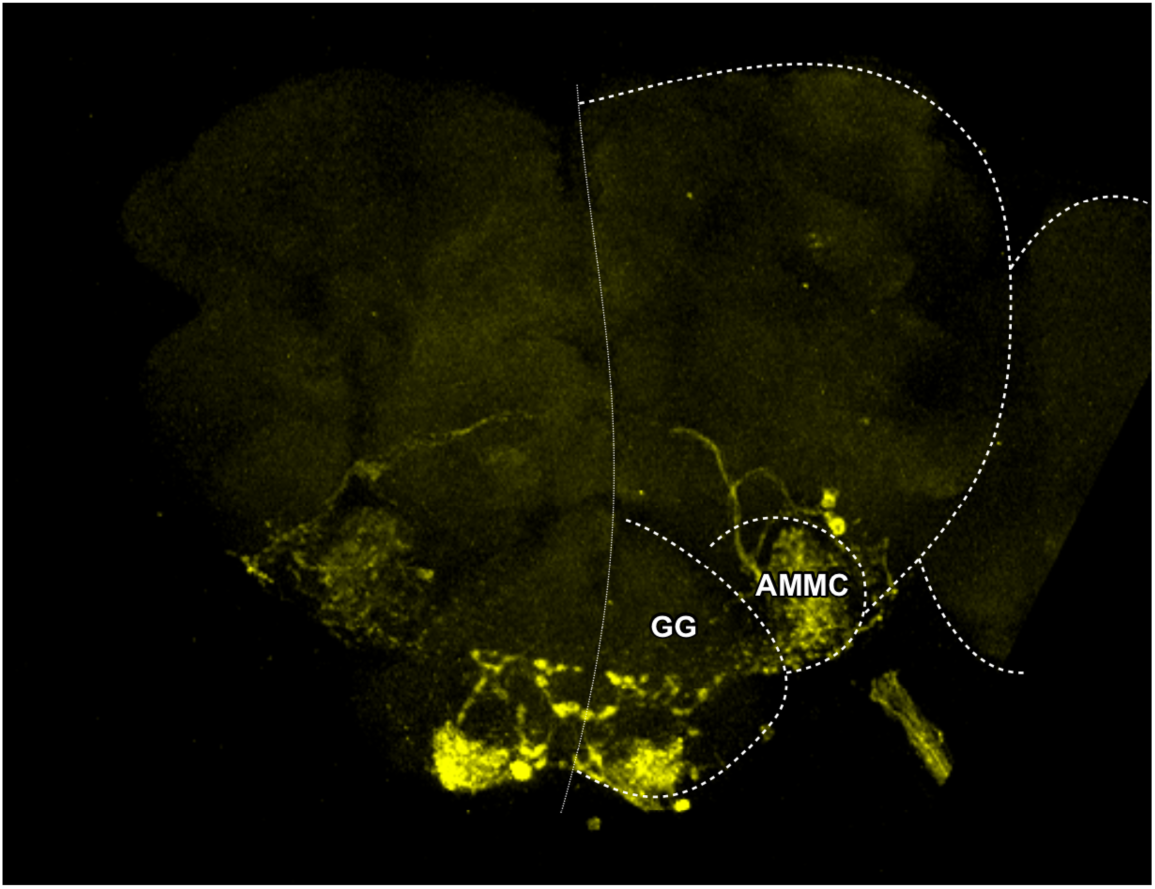
Staining in a *trans*-Tango control animal. Neurons in the gnathal ganglion (GG) and antennal mechanosensory and motor center (AMMC) stained positive in control animals of the *trans*-Tango/iso^KH11^ genotype (heterozygous for the *trans*-Tango transgenic label, but lacking a Gal4 to drive its expression). Iso^KH11^ is an inbred *w*^*1118*^ strain that was the common genetic background for our experiments, but otherwise harbors no genetic elements that could obviously explain this staining. Since the staining in this region was not specific to the P-F-R or PF-LC Gal4 drivers, we masked this region out of the reconstruction and analysis of PF-LCs postsynaptic partners. This staining was not present in P-F-R>*trans*-Tango animals. Dotted line indicates the midline; dashed lines outline neuropils.

**Figure S6.**
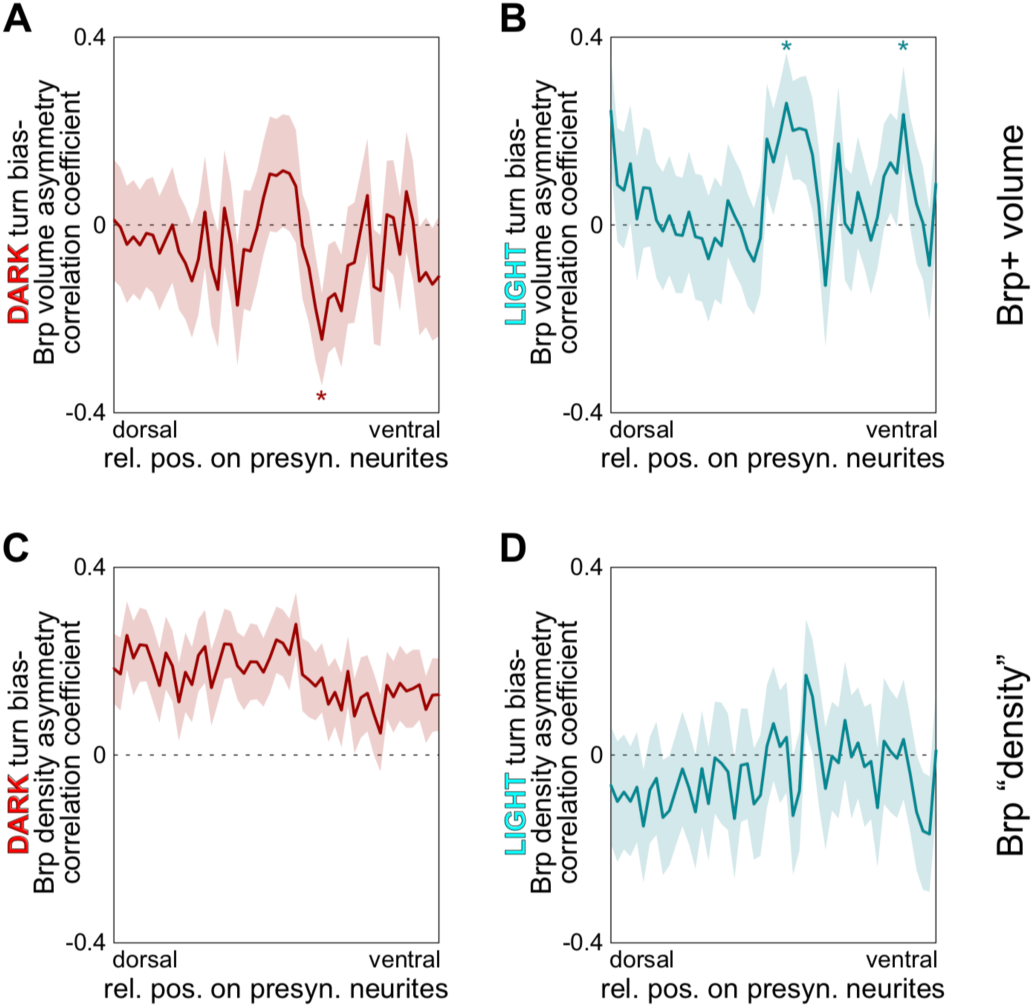
Controls for the correlation of PF-LC individual variation and turn bias behavior. **A**) Correlation between left-right asymmetry of the volume of Brp^short^-positive voxels in PF-LC LAL presynaptic compartments and turn bias exhibited in the dark, as a function of position along the long axis of the projections. Cyan shaded region is +/-SE of the correlation as determined by bootstrapping. **B**) As in A, but for the correlation between PF-LC LAL volume asymmetry and the turn bias exhibited in the light. **C, D**) As in A and B, but for Brp “density,” the ratio of Brp intensity (Fig 6E,F) to volume (A,B).

**Figure S7.**
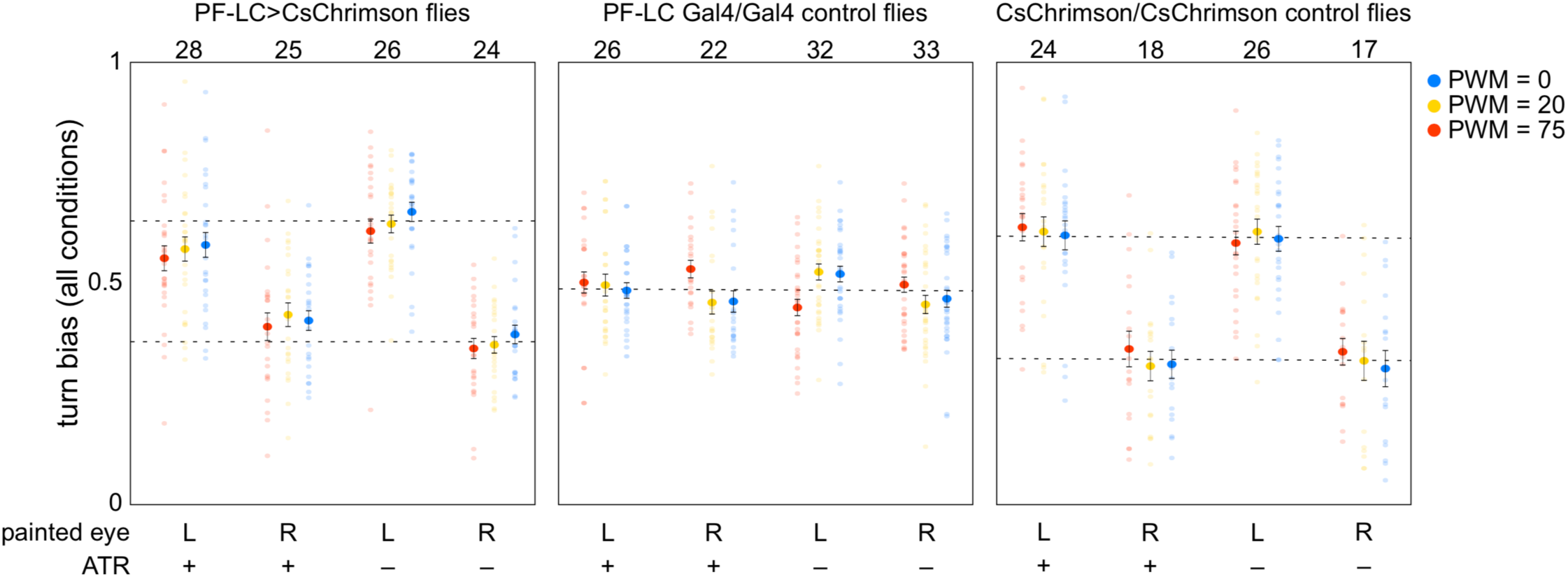
Turn biases of individual flies in the chronic PF-LC optogenetic experiment, by luminance. Turn bias of individual flies (points) with left or right eyes painted, with or without ATR, following 5 days of chronic illumination. Colors indicate the brightness of white LEDs during behavioral testing: pulse-width modulation (PWM) duty cycles of 0, 20 and 75/255. Large dots are the experimental group means, and bars +/-SEM. Numbers indicate sample sizes. Dashed lines indicate the approximate mean of the unpainted data points. Control genotypes (right two panels) do not show an optogenetic-specific change in turn bias.

**Movie S1.**
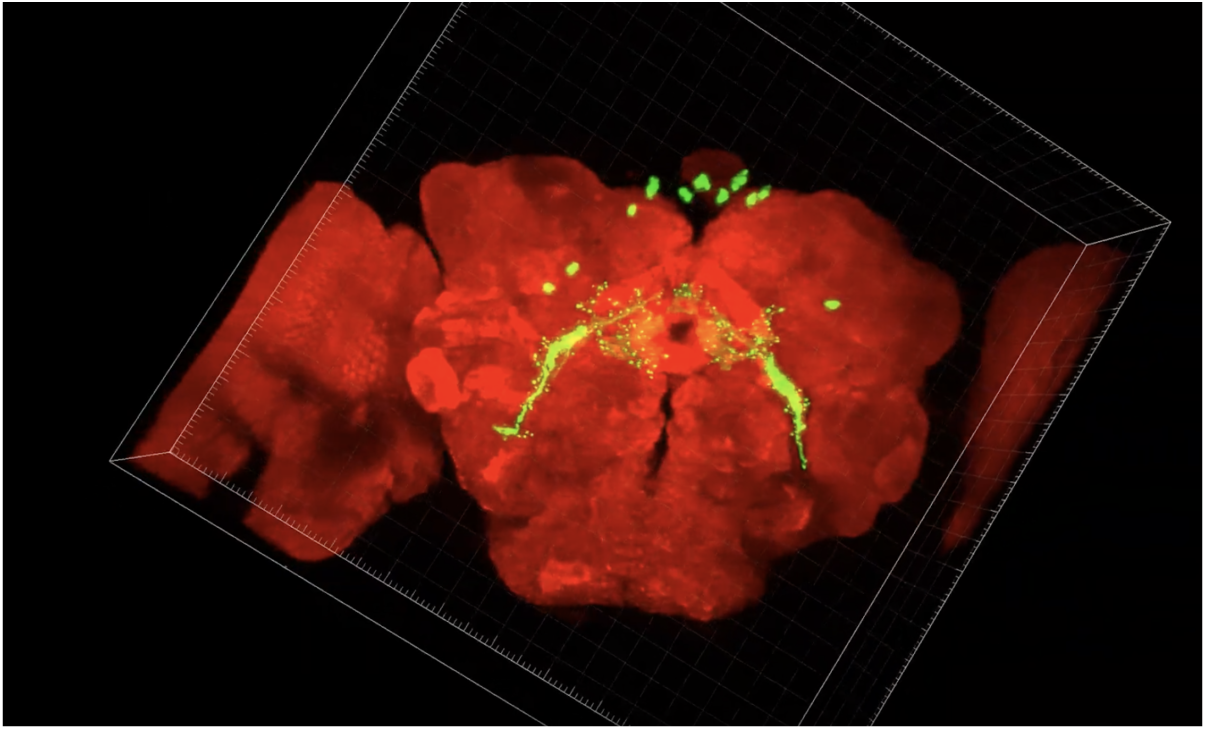
*https://youtu.be/TyCIsqpOef0* — Antibody staining of PF-LC pre-synapses. Red channel is an *α*-nc82 counterstain labeling synapses, green channel is *α*-Brp^short^.

**Movie S2.**
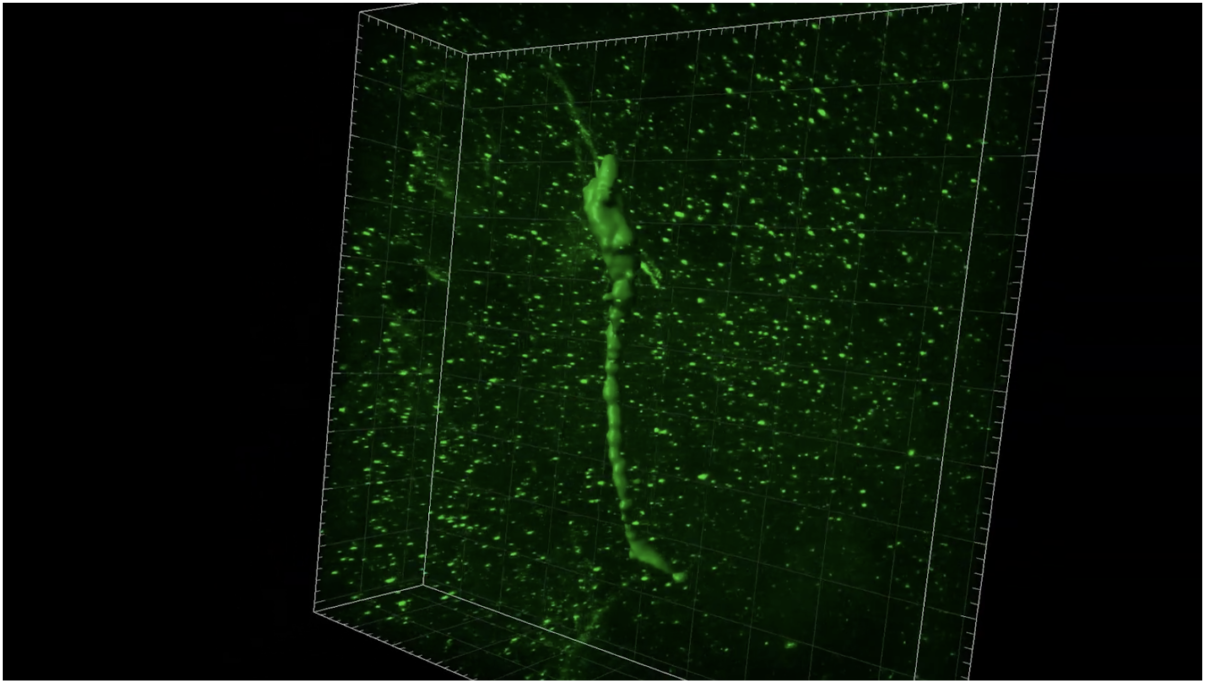
*https://youtu.be/iyQRXsRH46M* — Visualization of the erosion-dilation procedure for quantifying PF-LC axonal arbor anatomy. Stages of image processing consisted of creating surfaces in Imaris 9.0 that were increasingly tight to the neurite. Surfaces were created around adjacent voxels above a designated intensity threshold, and voxels outside of the surface are eroded away. Surface 1 approximates neurite and excludes noise. Surface 2 and 3 increasingly tighten to neurite to capture contours.

**S1 Table:**
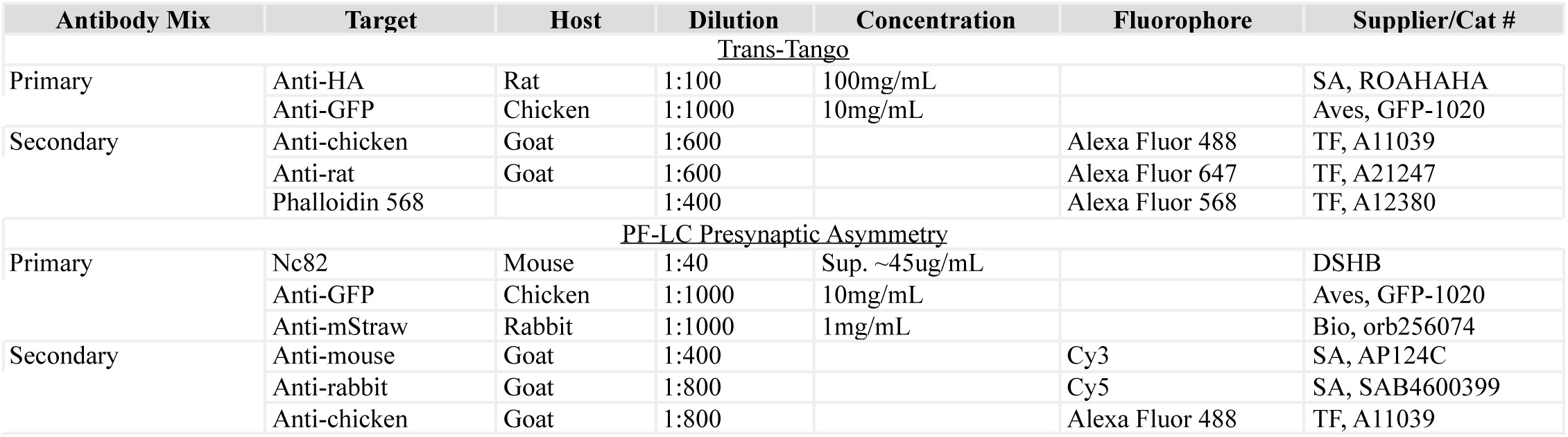
Antibodies in this study — All mixes diluted with 5% Goat Serum in PBT.

**S2 Table.** Strain table.

| Name | Allele | Source | Stock # | Reference | Target |
| --- | --- | --- | --- | --- | --- |
| SS02207-Gal4 |  | G. Rubin, T. Wolff |  | Wolff et al., 2018 | P-FNP |
| SS00425-Gal4 |  | G. Rubin, T. Wolff |  | Wolff et al., 2018 | P-FNP |
| SS00044-Gal4 |  | G. Rubin, T. Wolff |  | Wolff et al., 2018 | P-FNP |
| SS02221-Gal4 |  | G. Rubin, T. Wolff |  | Wolff et al., 2018 | P-FNP |
| SS02304-Gal4 |  | G. Rubin, T. Wolff |  | Wolff et al., 2018 | P-FNP |
| SS00418-Gal4 |  | G. Rubin, T. Wolff |  | Wolff et al., 2018 | P-FNP |
| SS00279-Gal4 |  | G. Rubin, T. Wolff |  | Wolff et al., 2018 | P-FNP |
| SS00079-Gal4 |  | G. Rubin, T. Wolff |  | Wolff et al., 2018 | P-FNA |
| SS00406-Gal4 |  | G. Rubin, T. Wolff |  | Wolff et al., 2018 | P-FNA |
| SS02255-Gal4 |  | G. Rubin, T. Wolff |  | Wolff et al., 2018 | P-FNA |
| SS02221-Gal4 |  | G. Rubin, T. Wolff |  | Wolff et al., 2018 | P-FNA |
| SS00007-Gal4 |  | G. Rubin, T. Wolff |  | Wolff et al., 2018 | P-FNA |
| SS02304-Gal4 |  | G. Rubin, T. Wolff |  | Wolff et al., 2018 | P-FNA |
| SS00083-Gal4 |  | G. Rubin, T. Wolff |  | Wolff et al., 2018 | P-FNA/D |
| SS00161-Gal4 |  | G. Rubin, T. Wolff |  | Wolff et al., 2018 | P-FND |
| SS00078-Gal4 |  | G. Rubin, T. Wolff |  | Wolff et al., 2018 | P-FND |
| SS00019-Gal4 |  | G. Rubin, T. Wolff |  | Wolff et al., 2018 | P-FND |
| SS00082-Gal4 |  | G. Rubin, T. Wolff |  | Wolff et al., 2018 | P-FND/V |
| SS02243-Gal4 |  | G. Rubin, T. Wolff |  | Wolff et al., 2018 | P-FNV |
| SS02209-Gal4 |  | G. Rubin, T. Wolff |  | Wolff et al., 2018 | P-FNV |
| SS02211-Gal4 |  | G. Rubin, T. Wolff |  | Wolff et al., 2018 | P-FNV |
| SS02241-Gal4 |  | G. Rubin, T. Wolff |  | Wolff et al., 2018 | P-EN |
| SS02268-Gal4 |  | G. Rubin, T. Wolff |  | Wolff et al., 2018 | P-EN |
| SS02232-Gal4 |  | G. Rubin, T. Wolff |  | Wolff et al., 2018 | P-EN |
| SS00097-Gal4 |  | G. Rubin, T. Wolff |  | Wolff et al., 2018 | P-EG |
| SS00090-Gal4 |  | G. Rubin, T. Wolff |  | Wolff et al., 2018 | E-PG |
| SS00131-Gal4 |  | G. Rubin, T. Wolff |  | Wolff et al., 2018 | E-PG |
| SS02258-Gal4 |  | G. Rubin, T. Wolff |  | Wolff et al., 2018 | E-PG |
| SS04781-Gal4 |  | G. Rubin, T. Wolff |  | Wolff et al., 2018 | E-PGT |
| SS04782-Gal4 |  | G. Rubin, T. Wolff |  | Wolff et al., 2018 | E-PGT |
| SS02374-Gal4 |  | G. Rubin, T. Wolff |  | Wolff et al., 2018 | E-PGT |
| SS02267-Gal4 |  | G. Rubin, T. Wolff |  | Wolff et al., 2018 | P-F-GS |
| SS02271-Gal4 |  | G. Rubin, T. Wolff |  | Wolff et al., 2018 | P-F-GS |
| SS02192-Gal4 |  | G. Rubin, T. Wolff |  | Wolff et al., 2018 | P-F-R |
| SS02293-Gal4 |  | G. Rubin, T. Wolff |  | Wolff et al., 2018 | P-F-R |
| SS02252-Gal4 |  | G. Rubin, T. Wolff |  | Wolff et al., 2018 | PF-LC |
| SS02214-Gal4 |  | G. Rubin, T. Wolff |  | Wolff et al., 2018 | PF-LC |
| SS02216-Gal4 |  | G. Rubin, T. Wolff |  | Wolff et al., 2018 | LPs-P |
| SS02200-Gal4 |  | G. Rubin, T. Wolff |  | Wolff et al., 2018 | LPs-P |
| SS04775-Gal4 |  | G. Rubin, T. Wolff |  | Wolff et al., 2018 | P6-8P9 |
| SS02231-Gal4 |  | G. Rubin, T. Wolff |  | Wolff et al., 2018 | Delta7 |
| SS02235-Gal4 |  | G. Rubin, T. Wolff |  | Wolff et al., 2018 | Delta7 |
| SS00116-Gal4 |  | G. Rubin, T. Wolff |  | Wolff et al., 2018 | Delta7 |
| SS02296-Gal4 |  | G. Rubin, T. Wolff |  | Wolff et al., 2018 | Sps-P |
| SS00155-Gal4 |  | G. Rubin, T. Wolff |  | Wolff et al., 2018 | Sps-P |
| SS00153-Gal4 |  | G. Rubin, T. Wolff |  | Wolff et al., 2018 | Sps-P |
| SS04769-Gal4 |  | G. Rubin, T. Wolff |  | Wolff et al., 2018 | IbSps-P |
| SS03951-Gal4 |  | G. Rubin, T. Wolff |  | Wolff et al., 2018 | IbSps-P |
| norpA | *norpAP24* | Bloomington | 9048 | Hardie, 2003 | R1-8 |
| Rhodopsin1 | *ninaE8* | B. Song, D. Rogulja | 1861 |  | R1-6 |
| eyes absent |  | B. Song, D. Rogulja | 2285 |  | eyes |
| GMR-hid |  | B. Song, D. Rogulja |  | Bergmann, 1998 | R1-8 |
| cryptochrome |  | B. Song, D. Rogulja |  |  | Cry+ cells |
| Rhodopsin1 | *ninaE7* | B. Song, D. Rogulja | 2103 |  | R1-6 |
| GMR-Gal4 |  | B. Song, D. Rogulja |  |  | R1-8 |
| Rh1-Gal4 |  | B. Song, D. Rogulja |  |  | R1-6 |
| Rh3-Gal4 |  | B. Song, D. Rogulja |  |  | Some R7 |
| Rh4-Gal4 |  | B. Song, D. Rogulja |  |  | Some R7 |
| Rh5-Gal4 |  | B. Song, D. Rogulja |  |  | Some R8 |
| Pan-R7-Gal4 |  | B. Song, D. Rogulja |  |  | R7 |
| Pan-R8-Gal4 |  | B. Song, D. Rogulja |  |  | R8 |
| CS |  |  |  |  | Wild type |
| w1118 | *w1118IsoKH11* | K. Honegger |  | Honegger & Smith et al., 2019 | Wild type |
| UAS-dTRPA1 | *P{UAS-TrpA1(B).K}attP16* | Bloomington | 26263 | Hamada, 2008 |  |
| UAS-Shibirets | *PBac{20XUAS-TTS-shi[ts1]-p10}attP2 l* | G. Rubin, Y. Aso |  |  |  |

